# An HWE/HisKA2-family kinase-phosphatase pair differentially regulates two paralogous PhyR response regulators to shape the General Stress Response specificity in *Sphingopyxis granuli* TFA

**DOI:** 10.64898/2026.11.12.766179

**Authors:** Alberto Pires-Acosta, Inmaculada García-Romero, Francisca Reyes-Ramírez

## Abstract

The general stress response (GSR) enables bacteria to withstand diverse environmental challenges. In Alphaproteobacteria, the GSR is controlled by a partner-switching mechanism involving the extracytoplasmic function sigma factor EcfG, its anti-sigma factor NepR, and the response regulator PhyR. Phosphorylated PhyR binds to NepR, releasing EcfG to activate GSR gene expression. However, how upstream signalling coordinates multiple PhyR paralogs remains poorly understood.

*Sphingopyxis granuli* TFA is a pollutant-degrading bacterium able to use the organic solvent tetralin as a sole carbon and energy source. TFA contains two copies of each core GSR regulatory component, including two PhyR paralogs with distinct contributions to GSR activity and stress resistance.

Using genetic, physiological and biochemical approaches, we identify two HWE/HisKA2-family proteins, HK_3483 and HK_1165, as major positive and negative GSR regulators, respectively. HK_3483 functions as a histidine kinase required for effective GSR activation through either PhyR paralog and for resistance to oxidative stress, copper and desiccation. It directly phosphorylates both PhyR proteins *in vitro*, with a preference for PhyR1. In contrast, HK_1165 preferentially dephosphorylates PhyR2. Consistently, deletion of *SGRAN_1165* in a *ΔphyR1* background increases PhyR2 phosphorylation *in vivo* under oxidative stress and stationary phase. Loss of HK_1165 causes a pronounced growth defect that is suppressed by deletion of either PhyR paralog while the remaining PhyR paralog supports elevated GSR activity and broad stress resistance.

Our findings support a model in which differential regulation of PhyR phosphorylation, mediated by the opposing activities of an upstream kinase-phosphatase pair, contributes to functional specialization within bacterial stress signalling networks.

## Introduction

Bacteria are continuously exposed to fluctuations in environmental conditions that challenge cellular homeostasis and survival. To cope with these changes, they rely on complex signalling pathways that detect environmental cues and coordinate adaptive cellular responses. In addition to dedicated stress response pathways that respond to specific insults, such as the SOS response to DNA damage (Maslowska et al., 2019), the heat shock response to protein misfolding (Richter et al., 2010), and the oxidative stress response (Ezraty et al., 2017), many bacteria possess a General Stress Response (GSR), a global regulatory system that provides broad protection against multiple and often unrelated environmental stresses (Gottesman, 2019).

The molecular mechanisms governing the GSR differ considerably among bacterial groups. In Gammaproteobacteria such as *Escherichia coli*, the GSR is controlled by the alternative sigma factor RpoS (σS), however, Alphaproteobacteria lack RpoS homologues (Staroń and Mascher, 2010). Instead, in many members of this class, the GSR is governed by a conserved partner-switching mechanism involving three core regulatory proteins: the extracytoplasmic function (ECF) sigma factor EcfG, its cognate anti-sigma factor NepR, and the anti-anti-sigma response regulator PhyR. Under non-stress conditions, NepR sequesters EcfG preventing its association with the RNA polymerase and thereby restricting transcription of GSR genes. Upon stress exposure, phosphorylation of a conserved aspartate residue within the receiver domain of PhyR induces a conformational change that exposes its N-terminal sigma-like domain, enabling PhyR to bind NepR with high affinity. This interaction releases EcfG, allowing its association with RNA polymerase and activation of the GSR regulon. Because the N-terminal domain of PhyR structurally and functionally mimics EcfG in its interaction with NepR, this regulatory strategy has been termed sigma-factor mimicry (Francez-Charlot et al., 2009; Herrou et al., 2012). Thus, control of PhyR phosphorylation is a central regulatory point controlling activation of the alphaproteobacterial GSR.

Although the molecular basis of the PhyR–NepR–EcfG partner-switching mechanism is now well established, considerably less is known about the upstream signalling pathways that control PhyR phosphorylation. Histidine kinases belonging to the HWE/HisKA2 family have emerged as major regulators of the alphaproteobacterial GSR. They constitute a distinct group of sensor kinases characterized by conserved sequence motifs in their catalytic domains that differ from those of canonical histidine kinases. Specifically, in their dimerization and histidine phosphotransfer (DHp) and catalytic ATP-binding domains (Herrou et al., 2017; Karniol and Vierstra, 2004). Unlike conventional two-component systems, in which a histidine kinase typically phosphorylates a cognate response regulator, HWE/HisKA2-family proteins often participate in more complex signalling networks involving multiple regulatory proteins and phosphotransfer steps. Such architectures provide opportunities for integrating diverse environmental signals and fine-tuning cellular responses (Chen et al., 2025; Fiebig et al., 2019; Foreman et al., 2012; Kaczmarczyk et al., 2015; Kim et al., 2014; Lori et al., 2018; Wülser et al., 2022).

The mechanisms connecting HWE/HisKA2 kinases to PhyR differ considerably among alphaproteobacterial species. In *Brucella abortus*, the HWE/HisKA2 kinase LovhK specifically phosphorylates PhyR *in vitro* (Kim et al., 2014). By contrast, in *Sphingomonas melonis* Fr1, multiple HWE/HisKA2 histidine kinases, termed PhyR-activating kinases (Paks), respond to partially overlapping sets of stresses. Upon stress exposure, phosphoryl groups are transferred sequentially from the Paks to the single-domain response regulator (SDRR), SdrG, then to PhyT, a membrane-associated phosphotransfer protein and ultimately to PhyR, promoting its activation and triggering the partner-switching mechanism (Gottschlich et al., 2018; Kaczmarczyk et al., 2014). Similarly, in *Caulobacter crescentus*, multiple HWE/HisKA2 kinases activate PhyR through a phosphorelay involving the SDRR MrrA and the histidine phosphotransferase PhyK (Lori et al., 2018).

Beyond the activation of PhyR, mechanisms that limit its phosphorylation are also important to prevent inappropriate GSR activation. However, comparatively few examples have been characterized in Alphaproteobacteria. In *Sphingomonas melonis* Fr1, PhyT also contributes to restricting PhyR activation under non-stress conditions through complex formation with unphosphorylated PhyR, (Gottschlich et al., 2018). In *Sinorhizobium meliloti,* genetic evidence indicates that the HWE-family histidine kinase RsiC exerts both positive and negative control over the GSR pathway involving the two functionally redundant PhyR homologues, RsiB1 and RsiB2 (Sauviac and Bruand, 2014). Despite these examples, the biochemical mechanisms underlying negative regulation of PhyR phosphorylation remain poorly characterized.

*Sphingopyxis granuli* TFA is an environmental alphaproteobacterium originally isolated from Rhine River sediment and capable of using the organic solvent tetralin as its sole carbon and energy source (Hernáez et al., 1999). It is the only member of its genus described as a facultative anaerobe, being able to grow under anoxic conditions using nitrate as a terminal electron acceptor (González-Flores et al., 2019). This metabolic versatility may facilitate its adaptation to environments with fluctuating oxygen availability and highlights its potential relevance for the bioremediation of organic pollutants. A distinctive feature of the GSR system in TFA is the duplication of all three core regulatory components, including two EcfG sigma factors (EcfG1 and EcfG2), two anti-sigma factors (NepR1 and NepR2), and two response regulators (PhyR1 and PhyR2). Using a combination of genetic and biochemical studies, including *in vitro* transcription assays, we previously demonstrated that these duplicated regulators are not functionally redundant but instead perform distinct roles in controlling GSR activation. In particular, PhyR1 and PhyR2 contribute differently to resistance against different environmental stresses and generate different levels of GSR activation during stationary phase, revealing functional specialization within the signalling cascade (De Dios et al., 2022). However, the upstream signalling mechanisms that differentially regulate the activity of each PhyR paralog remain unknown.

Here, we investigate the contribution of HWE/HisKA2-family histidine kinases to GSR regulation in *S. granuli* TFA. Using genetic, physiological, and biochemical approaches, we identify HK_3483 and HK_1165 as major positive and negative regulators of the pathway, respectively. We demonstrate that HK_3483 directly phosphorylates both PhyR paralogs, *in vitro*, whereas HK_1165 exhibits phosphatase activity primarily toward PhyR2. Our results reveal that differential control of PhyR phosphorylation modulates signalling through the two PhyR paralogs and provides a mechanism for generating distinct GSR outputs. These results offer new insights into how phosphorylation and dephosphorylation shape signalling specificity in the GSR of pollutant-degrading alphaproteobacteria.

## Materials and Methods

### Media growth and conditions

*Escherichia coli* was grown in Luria-Bertani (LB) medium. at 37°C. *Sphingopyxis granuli* strains were grown at 30 °C in MML mineral medium. Where indicated, MM minimal medium supplemented with β-hydroxybutyrate (BHB) at a final concentration of 8mM was used to grow *S. granuli* TFA. When required, solid and liquid media were supplemented with kanamycin, ampicillin or streptomycin at the concentrations indicated in Supplementary Table S1.

### Construction of strains and plasmids

Strains, plasmids, and oligonucleotides used in this work can be found in Supplementary material. In-frame deletions in *SGRAN_1165*, *SGRAN_1773*, *SGRAN_2544* and *SGRAN_3483* were constructed as described in (García-Romero et al., 2024). Upstream and downstream flanking regions of the target gene were amplified with specific oligonucleotides (Suplementary Table S3). Resulting PCR fragments were digested with corresponding restriction enzymes and cloned into pMPO1412 vector digested with compatible enzymes. After verification by PCR and sequencing for validation the resulting vectors (Suplementary Table S2) were introduced into competent TFA cells via electroporation.

Transformants were selected on agar plates supplemented with 20 mg/L of kanamycin (Km) and then screened for sensitivity to 200 mg/L of streptomycin (Str). Plasmid integration into the genome was validated by PCR amplification of the kanamycin resistance cassette using the oligonucleotides KmFw-pk18 and KmRev-pK18.

To force a second recombination event, pSW-I vector, which expresses the I-SceI endonuclease, was introduced by electroporation. Basal expression of the endonuclease was sufficient to cut the I-SceI sites, so no induction was required. Candidates that had undergone the second recombination were identified by their loss of kanamycin resistance. Gene deletion was confirmed by PCR screening of the candidate colonies using specific oligonucleotides. Finally, the pSW-I vector was cured from the mutant strain by repeated subculturing in the absence of ampicillin.

Successful loss of the plasmid was confirmed by the resulting loss of ampicillin resistance. Final validation of the deletion was achieved by sequencing the PCR products.

Strains expressing 3×FLAG-tagged PhyR2 for immunoblot detection were generated by the same allelic exchange procedure using pMPO1460 (De Dios et al., 2022).

### Genome sequencing to identify suppressor mutations

*ΔSGRAN_1165* strains obtained independently and suspected of carrying suppressor mutations were sequenced by MicrobesNG (Birmingham, UK) using Illumina paired-end technology. Reads were mapped against the complete genome of *Sphingopyxis granuli* TFA (GenBank accession no. CP012199), used as the wild type reference. Single-nucleotide polymorphisms and small insertions/deletions were identified with breseq (Deatherage and Barrick, 2014) which uses Bowtie2 for read alignment (Langmead and Salzberg, 2012) The mutations detected were annotated with gdtools ANNOTATE (Deatherage and Barrick, 2014).

### Stress resistance assays

Assays to test the resistance to different stressors were performed as described by (De Dios et al., 2020) with some modifications. For osmotic and heavy metal stress, late-exponential phase cultures were adjusted to an OD 600 of 1 and serially diluted, 10µl were then placed on were placed into MML agar plates supplemented with either 550mM of NaCl or 3.5mM of CuSO_4_. Plates were incubated at 30°C during 5-6 days until biomass was observed.

For desiccation assays, 5 μl spots of serial dilutions were placed on 0.45 μm pore size filters (Sartorius Stedim Biotech GmbH) and left to air dry in a laminar flow cabin to dry for 2 hours (with control plates only dried for 5 minutes) and kept at room temperature for an additional 20 hours. Then, filters were placed in MML supplemented with bromophenol blue 0.002% and incubated for 5-6 days at 30°C.

For oxidative stress assays, late-exponential phase cultures were diluted to an OD 600 of 0.1, and cultures were allowed to grow to an OD 600 of 0.5. On that moment, H_2_O_2_ was added to a final concentration of 10mM. Recovery from oxidative shock was calculated by comparing OD 600 nm after 3 hours of growth in treated and untreated cultures with results represented as a percentage.

### β-galactosidase activity measurement

As a reporter of GSR cascade activation, a *nepR2::lacZ* translational fusion was integrated into the chromosome of different TFA strains using the pMPO1408 vector (De Dios et al., 2020). Saturated precultures grown in (MM) minimal medium were diluted to an OD 600 of 0.1 in MM supplemented with 8mM β-hydroxybutyrate. Samples were taken at 10 hours for exponential phase and at 48 hours for stationary phase, and β-galactosidase activity was measured as described by (Miller, 1972). Data are presented as the means of three independent biological replicates.

### Phosphorylation state detection by Phos-tag^tm^ SDS-PAGE

*3xflag* fusion in either *phyR1* or *phyR2* were introduced in different TFA strains. For oxidative stress assays, cultures were grown to an OD₆₀₀ of 0.5 and treated with 10 mM H_2_O_2_. Samples were collected 3 h after H_2_O_2_ addition. For stationary-phase analysis, samples were collected separately from stationary-phase cultures, after 48 h of growth in MM supplemented with 8 mM BHB.

Pellets were resuspended in Tris-HCl buffer (pH 8). Total protein concentration was quantified by the Bicinchoninic Acid (BCA) method (Smith et al., 1985) and 10µg of protein were finally loaded into 12% SDS-polyacrylamide gels supplemented with 25 μM Phos-tag^tm^ acrylamide (NARD Chemicals, Hiroshima, Japan) and 50 μM MnCl2 (Kinoshita et al., 2006). Samples were not boiled before loading as high temperatures might break aspartate-phosphate bonds (Barbieri and Stock, 2008).

Electrophoresis was conducted at 80V and at 4°C and gels were incubated with transfer buffer supplemented with 10 mM EDTA to chelate Mn^2+^ ions and improve transfer efficiency. After that, proteins were transfer to nitrocellulose membrane using a Trans-Blot® Turbo^TM^ semi-dry transfer system (Bio-Radand). Finally, proteins were detected using anti-flag mouse primary antibodies (Sigma Aldrich) and anti-mouse secondary antibodies conjugated to peroxidase (Sigma Aldrich).

### Protein overexpression and purification

Full-length HK_1165 and truncated variant of HK_3483 proteins were purified using the IMPACT system (New England Biolabs) using manufacturer’s instructions. pTYB21_3483 and pTYB21_1165 vectors (containing the Histidine kinase core of HK_3483 and the whole HK_1165 gene respectively) were transformed into expression strain *E. coli* ER2566. OD 600 of 0.1 LB cultures was prepared from saturated preinocula in a total volume of 400ml. Cultures were incubated at 37°C 180 rpm until they reached OD 600 0.5. At that point, they were chilled in ice for 30 minutes, induced with 0.5mM IPTG and incubated at 20°C (overnight in the case of HK_3483 and for three hours for HK_1165).

After incubation, cells were harvested, resuspended in 20 ml of column buffer, and lysed by sonication. The clarified lysates were loaded in a chitin resin column left to flow through the resin by gravity at a low flow rate. After that, column was flushed with 10 volumes of column buffer and then with 5 volumes of column buffer supplemented 10mM of DTT to release the protein from the intein. Columns were incubated with the lysis buffer at 4°C for 48 h approximately. The presence of the target protein was assessed by SDS-PAGE. Then, DTT concentration in the buffer was reduced by dialysis against kinase buffer (5mM DTT, 300mM HEPES-KOH pH8, 50mM de KCl mM 5mM MgCl_2_) at 4°C overnight using a 3 kDa pore size dialysis cassette (Thermo Fisher Scientific). Purity and concentration of the protein mixtures were evaluated by densitometry compared to different dilutions of BSA using a Typhoon scanner and the ImageLab software. Finally, protein mixtures were aliquoted and frozen at −80°C.

### *In vitro* phosphorylation assays

HK_3483 autophosphorylation assays were performed with the purified kinase domain of HK_3483. 100µl mix of 10µM HK_3483 in kinase buffer was prepared and reactions were started by adding a mix of 1mM cold ATP and 50µCi of radiolabeled ATP, [γ-32P]-6000Ci/mmol to the reaction probe. Reactions were stopped at different time intervals by taking 10µl samples and adding 10µl of 2xloading buffer 160 mM Tris–HCl, pH 8.0, 20% [v/v] glycerol, 4% [w/v] SDS, and 0.1% [w/v] bromophenol blue) to stop the reaction.

Samples were separated in a 12.5% acrylamide gel and ran at 120V for 40 minutes after which gels were dried, the dye front removed and exposed to a phosphorimager screen overnight at -80°C. Images were obtained by scanning the screens in a Typhoon 9410 scanner and relative intensity was later quantified with ImageJ.

For phosphotransfer reactions to either PhyR1 or PhyR2, we prepared again a 10µM HK_3483 mix in kinase buffer and incubated for 1 hour after adding ATP to obtain phosphorylated HK_3483. Excess ATP was removed with a Zeba desalting column MW 7000 Da. Equal volumes of HK_3483 mix and a 10µM PhyR1 or PhyR2 mix were put together to a final concentration of 5µM for both proteins initiating phosphotransfer reaction. Once again, 10µl samples were taken at different time intervals and reactions were stopped by adding 10µl of 2xloading buffer as described above.

### *In vitro* dephosphorylation assays

For assessing the phosphatase activity of HK_1165 *in vitro* we prepared a 10µM HK_3483, 10µM PhyR1 or PhyR2 mix in kinase buffer and incubated for 1 hour to obtain phosphorylated PhyR1 and PhyR2. Excess ATP was removed with a Zeba desalting column MW 7000 Da and the desalted mix was incubated for 20 minutes more at room temperature. Equal volumes of HK_3483/PhyR mix and a 10µM HK_1165 mix were put together to a final concentration of 5µM for both proteins initiating phosphatase reaction. Once again, 10µl samples were taken at different time intervals and reactions were stopped by adding 10µl of 2xloading Buffer. Samples were analyzed by SDS–PAGE and phosphorimaging as described above.

## Results

### Identification and structural organization of candidate HWE/HisKA2 histidine kinases involved in GSR signaling

HWE/HisKA2 family histidine kinases constitute a distinct superfamily of sensor kinases with diverse roles in environmental signalling (Herrou et al., 2017; Karniol and Vierstra, 2004). As in other histidine kinases, their kinase core comprises a dimerization and histidine phosphotransfer (DHp) domain and a catalytic ATP-binding (CA) domain. However, HWE/HisKA2 kinases differ from canonical HisKA kinases in several features, including distinctive sequence motifs in both the catalytic and non-catalytic regions and in their catalytic machinery. A hallmark of this superfamily is the HRxxN motif within the DHp domain, which contains the phosphoryl-accepting histidine and provides a readily recognizable sequence signature of HWE/HisKA2 kinases. An additional characteristic is a conserved arginine in the second helix of the DHp domain, referred to as the R-box. The CA domain also displays distinctive features including the absence of the F-box and a short ATP lid. HWE/HisKA2 kinases can adopt different oligomeric states rather than the predominantly dimeric organization characteristic of canonical histidine kinases (Herrou et al., 2017).

In contrast to classical two-component systems, which are generally organized as cognate histidine kinase-response regulator pairs, HWE/HisKA2 kinases involved in the alphaproteobacterial GSR can form part of more complex signaling networks including those that channel environmental information through to the PhyR response regulator, to coordinate appropriate cellular responses (Fiebig et al., 2015; Kaczmarczyk et al., 2015; Kim et al., 2014; Wülser et al., 2022).

Using the characteristic HRxxN signature, analysis of the *S. granuli* TFA genome identified four candidate HWE/HisKA2 histidine kinases: HK_1165 (SGRAN_1165), HK_1773 (SGRAN_1773), HK_2544 (SGRAN_2544) and HK_3483 (SGRAN_3483). Based on their conserved kinase domains, HK_1165 and HK_3483 were classified as HWE-family kinases, whereas HK_1773 and HK_2544 belonged to the HisKA2 family. Their predicted domain organization and an alignment of conserved motifs within the DHp and CA domains are shown in Fig. 1. Although all four proteins share a conserved C-terminal kinase core, their predicted sensory domains and subcellular localizations differ. HK_3483 is the only predicted membrane-associated kinase and contains a predicted periplasmic CHASE sensory domain flanked by two transmembrane helices. HK_1165 and HK_2544 are predicted to be soluble cytoplasmic proteins containing GAF-PAS and PAS sensory domains, respectively. HK_1773 displays a distinct hybrid architecture, with an N-terminal receiver (REC) domain fused to the histidine kinase core (Fig. 1a).

**Figure 1.**
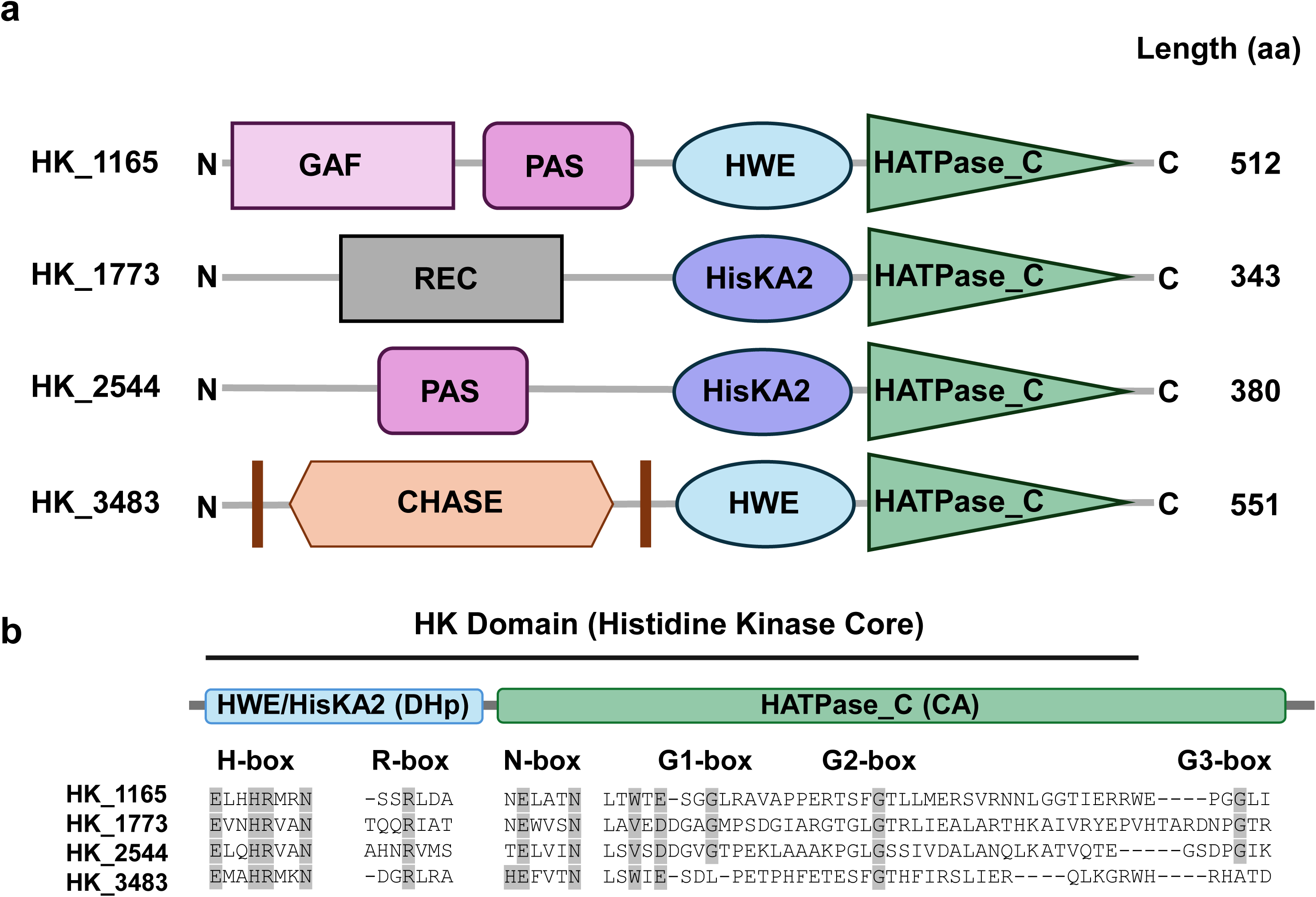
Domain organization and conserved sequence features of the four candidate HWE/HisKA2 histidine kinases of *Sphingopyxis granuli TFA*. (a) Schematic representation of the predicted domain organization of SGRAN_1165 (HK_1165), SGRAN_1773 (HK_1773), SGRAN_2544 (HK_2544), and SGRAN_3483 (HK_3483), as determined using InterPro (PMID: 39565202). Predicted sensory and regulatory domains, transmembrane helices, and the conserved histidine kinase core are indicated. (b) Sequence alignment showing amino acid residues corresponding to the conserved sequence motifs described by (Herrou et al., 2017). The H-box and R-box within the DHp domain and the N-box, G1-box, G2-box and G3-box within the CA domain are indicated. The characteristic HRxxN motif within the H-box includes the predicted phosphoryl-accepting histidine. Abbreviations: CHASE, cyclases/histidine kinases associated sensory extracellular; GAF, cGMP-specific phosphodiesterases, adenylyl cyclases and FhlA; PAS, Per-ARNT-Sim; REC, receiver domain; DHp, dimerization and histidine phosphotransfer; CA, catalytic ATP-binding domain

Given the involvement of HWE/HisKA2 kinases in regulating PhyR-dependent signaling in Alphaproteobacteria, these proteins were selected as candidate upstream regulators of the TFA GSR.

### The histidine kinase HK_3483 is a major positive regulator of GSR activation in *S. granuli* TFA

To investigate the contribution of these HWE/HisKA2 kinases to GSR regulation, we constructed single in frame-deletion mutants of the *SGRAN_1773*, *SGRAN_2544*, *SGRAN_3483* and *SGRAN_1165* genes. The phenotype associated with *ΔSGRAN_1165* is described separately below due to its distinct regulatory behavior. In addition, all possible double and triple mutant combinations of *SGRAN_1773*, *SGRAN_2544*, and *SGRAN_3483* were generated.

As previously reported, entry into stationary phase constitutes a GSR-inducing condition in *S*. *granuli* TFA (De Dios et al., 2020). To assess the contribution of the different HWE/HisKA2 kinases to GSR-dependent gene expression, a *nepR2::lacZ* translational fusion, previously validated as a reporter of GSR activity in TFA, was integrated into the chromosome of all kinase mutant backgrounds. The wild type strain and the *ΔphyR1ΔphyR2* mutant carrying the same fusion were included as controls. The *ΔphyR1ΔphyR2* mutant lacks both anti-anti-sigma factors is a GSR deficient strain, unable to activate the GSR and have a survival defect under several stress conditions (De Dios et al., 2020). β-galactosidase activity was quantified during exponential growth (non-stress condition), when the GSR remains largely repressed and stationary phase when the response is activated (stress condition).

As shown in Fig. 2a, during exponential growth, *nepR2::lacZ* activity remained low in the wild type and in the kinase mutants, indicating that these deletions did not affect the basal GSR activity. However, during stationary phase, only those strains lacking HK_3483, failed to induce the GSR response efficiently, since a marked fourfold reduction in *nepR2* promoter activity was observed with levels of *nepR2::lacZ* activity comparable to those of the wild type under non stress conditions. Mutants lacking HK_1773 and/or HK_2544 showed stationary-phase induction comparable to the wild type strain.

**Figure 2:**
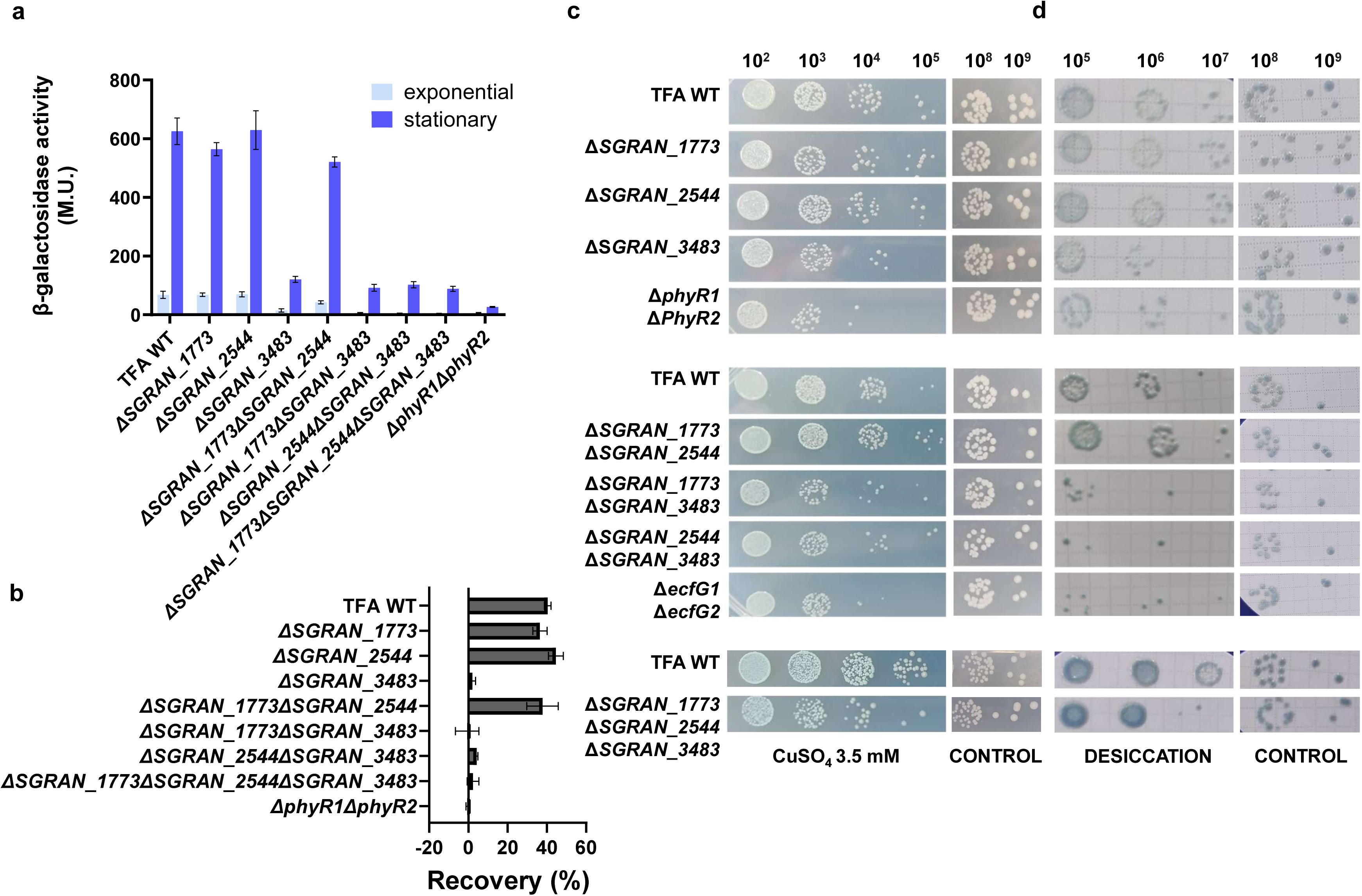
GSR expression and stress-resistance phenotypes of HWE/HisKA2 histidine kinase deletion mutants. GSR-dependent gene expression and stress resistance in single deletion mutants of *SGRAN_1773*, *SGRAN_2544*, and *SGRAN_3483*, and in double and triple mutant combinations. Wild-type TFA, and either the *ΔphyR1ΔphyR2* or *ΔecfG1 ΔecfG2* GSR-deficient strain were included as controls. (a) GSR-dependent gene expression assessed during exponential growth (10 h; non-stress condition) and stationary phase (48 h; GSR-inducing condition) in minimal medium supplemented with 8 mM β-hydroxybutyrate as the carbon source. Strains carrying a chromosomal *nepR2::lacZ* translational fusion were used, and reporter activity was determined by β-galactosidase assays.(b) Recovery from oxidative shock was determined by comparing the OD 600 of cultures treated with 10 mM H_2_O_2_ to untreated controls 3 h after exposure.(c) Copper stress sensitivity was assessed by spotting serial dilutions onto MML agar supplemented with 3.5 mM CuSO₄ and (d) Desiccation resistance was assessed by spotting serial dilutions onto filters, air-drying them for 20 h, and subsequently placing the filters onto MML agar supplemented with 0.002% bromophenol blue. Filters transferred directly onto the plates without air-drying were used as controls.

We next examined whether the impaired GSR activation associated with the absence of HK_3483 was accompanied by reduced resistance to various environmental stresses. The mutant strains were subjected to a series of selected stress conditions for which PhyR-dependent stress resistance has previously been demonstrated in TFA (De Dios et al., 2022, 2020), including oxidative stress, copper, and desiccation (Fig. 2b-d). As a control either *ΔecfG1ΔecfG2* double mutant lacking the two sigma factors or *ΔphyR1ΔphyR2* lacking both anti-anti-sigma factors of the GSR were included as indicated in each panel. This GSR deficient strain are unable to activate the GSR and have a survival defect under stress conditions.

Deletion of either, *SGRAN_1773* and *SGRAN_2544* did not result in differences in stress survival compared to the wild type strain under any of the stresses tested. In contrast, the *ΔSGRAN_3483* mutant displayed increased sensitivity to oxidative stress copper and desiccation, resembling the phenotype of the GSR-deficient control (Fig. 2b-d). Consistently, all double and triple mutants carrying the *SGRAN_3483* deletion exhibited similar sensitivity to the stresses as the *ΔSGRAN_3483* single mutant, while the *ΔSGRAN_1773ΔSGRAN_2544* double mutant behaved like the wild type strain (Fig. 2b-d). Thus, the defect in GSR activation observed in the absence of HK_3483 is accompanied by reduced stress resistance. These results identify HK_3483 as the major HWE/HisKA2 kinase required for effective GSR activation and resistance to the stress conditions tested.

The broad stress-sensitivity of the *ΔSGRAN_3483* mutant suggests that HK_3483 may sense different environmental signals acting as a major upstream activator of the GSR cascade in TFA. Mechanistically, this likely reflects its ability to promote PhyR phosphorylation, thereby enabling the partner-switching cascade that releases the EcfG sigma factors from NepR inhibition. In the absence of HK_3483, impaired PhyR activation would prevent proper induction of GSR-regulated genes, ultimately leading to an increased stress sensitivity.

To further determine that HK_3483 is required for signaling through both PhyR paralogs, double mutants combining the *SGRAN_3483* deletion with either *ΔphyR1* or *ΔphyR2* were constructed, and their stress resistance and GSR expression phenotypes were compared with the corresponding single *ΔphyR* mutants (Fig. 3a). As previously reported, PhyR1 and PhyR2 exhibit a certain degree of functional specialization in their contribution to stress resistance in TFA, with their contribution depending on the specific stress condition. The *ΔphyR1* mutant retains resistance to copper stress, suggesting that PhyR2 is sufficient to mediate resistance under this condition, whereas the *ΔphyR2* mutant remains resistant to oxidative stress indicating that PhyR1 can sustain the response to this stress (De Dios et al., 2022); and Fig. 3a and Fig. 3c respectively). These phenotypes provide suitable conditions to examine signalling through PhyR2 or PhyR1, respectively.

**Figure 3.**
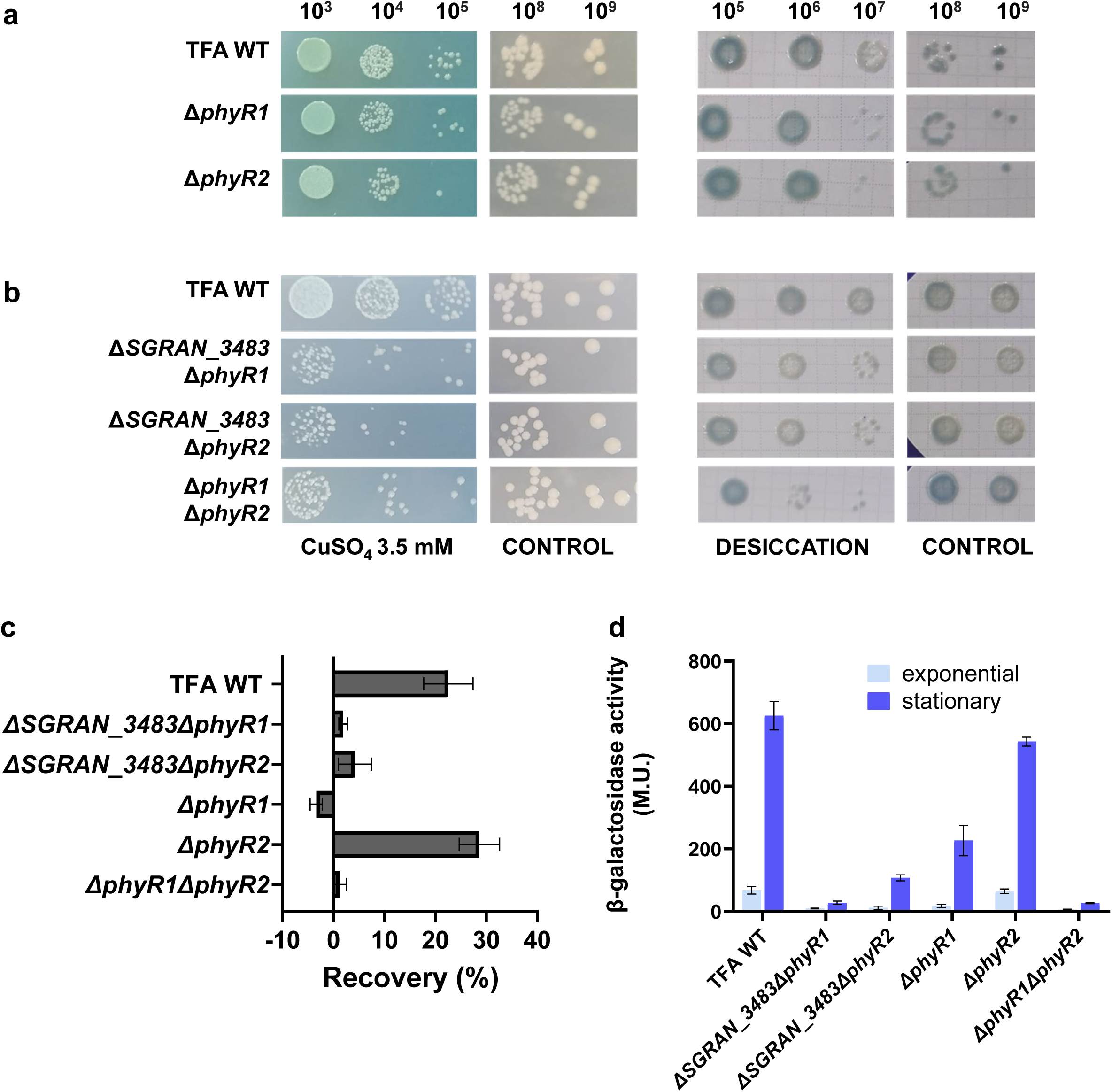
Stress-resistance and GSR expression phenotypes of *ΔSGRAN_3483ΔphyR1* and *ΔSGRAN_3483ΔphyR2 double mutants.* Stress resistance and GSR-dependent gene expression were assessed in the *ΔSGRAN_3483ΔphyR1* and *ΔSGRAN_3483ΔphyR2* double mutants and compared with wild-type TFA, the corresponding *ΔphyR1* and *ΔphyR2* single mutants, and the GSR-deficient *ΔphyR1ΔphyR2* double mutant. (a, b) Copper and desiccation stress resistance of the *ΔphyR1*, *ΔphyR2*, and *ΔphyR1ΔphyR2* mutants (a) and the *ΔSGRAN_3483ΔphyR1* and *ΔSGRAN_3483ΔphyR2* double mutants (b). Copper stress sensitivity was assessed by spotting serial dilutions onto MML agar supplemented with 3.5 mM CuSO₄. Desiccation resistance was assessed by spotting serial dilutions onto filters, air-drying them for 20 h, and subsequently placing the filters onto MML agar supplemented with 0.002% bromophenol blue. Filters transferred directly onto the plates without air-drying were used as controls. (c) Recovery from oxidative shock was assessed by comparing the OD 600 of cultures exposed to 10 mM H₂O₂ with that of untreated controls 3 h after treatment. (d) GSR-dependent gene expression was assessed during exponential growth (10 h; non-stress condition) and stationary phase (48 h; GSR-inducing condition) in minimal medium supplemented with 8 mM β-hydroxybutyrate as the carbon source. Strains carrying a chromosomal *nepR2::lacZ* translational fusion were used, and reporter activity was determined by β-galactosidase assays.

As shown in Fig. 3b, the *ΔSGRAN_3483ΔphyR1* mutant, which retains PhyR2 as the only paralog, displayed increased sensitivity to copper stress, compared with the *ΔphyR1* single mutant. Conversely, the *ΔSGRAN_3483ΔphyR2* mutant, which retains PhyR1, exhibited sensitivity to oxidative stress compared with the *ΔphyR2* single mutant. Both double mutants were also sensitive to desiccation (Fig 3b), although this phenotype was expected given the previously established requirement for both PhyR paralogs for full resistance to this stress (De Dios et al., 2022).

Consistent with their stress-sensitivity phenotypes, both *ΔSGRAN_3483ΔphyR1* and *ΔSGRAN3483ΔphyR2* are clearly impaired in GSR activation as determined using the *nepR2::lacZ* reporter (Fig. 3d). Thus, removal of HK_3483 compromises GSR activation regardless of which PhyR paralog remains in the cell.

These data genetically confirm that HK_3483 is a major upstream positive regulator required for effective signaling through both PhyR1 and PhyR2 under the conditions tested.

### HK_1165 negatively regulates PhyR-dependent GSR signaling

Among the four candidate HWE/HisKA2 histidine kinases identified in TFA, HK_1165 is the only one genetically associated with the GSR regulatory genes (Fig 4a) whereas the other three candidate kinases are encoded as orphan histidine kinases elsewhere in the genome. To assess the contribution of HK_1165 to GSR regulation, an in-frame deletion mutant was constructed using the two-step homologous recombination strategy optimized for TFA as employed for all strains constructed in this study. In this method, the cointegrate resulting from the first recombination event is resolve through a second recombination step either producing the deletion of the desired target gene or a restoration of the wild-type genotype, as detail in Material and Methods.

**Figure 4.**
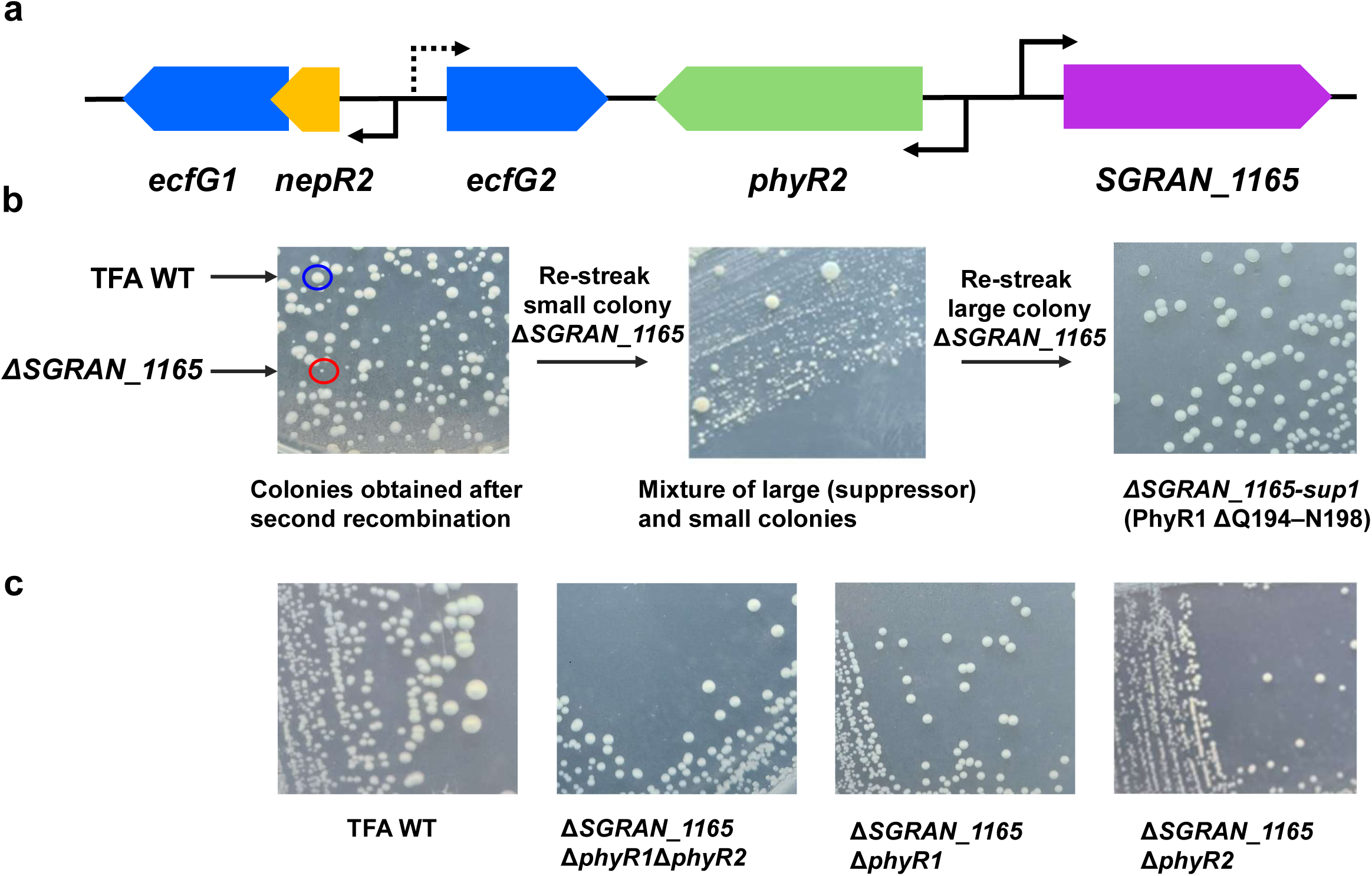
Suppression of the growth defect caused by loss of HK_1165. (a) Genetic organization of the GSR regulatory locus containing *SGRAN_1165* in *S. granuli* TFA. Genes encoding the EcfG sigma factors, NepR anti-sigma factors, PhyR response regulators, and HK_1165 are shown. Gene arrows indicate the direction of transcription. Solid arrows indicate transcriptional units belonging to the GSR regulon, whereas the dashed arrow represents *ecfG2*, which is not part of the GSR regulon. (b) Colony phenotype associated with deletion of *SGRAN_1165*. Following the second recombination event during construction of the *ΔSGRAN_1165* mutant, two distinct colony sizes were obtained. Large colonies (blue circle) corresponded to restoration of the wild-type allele, whereas small colonies (red circle) carried the *ΔSGRAN_1165* allele. Re-streaking of a small *ΔSGRAN_1165* colony on MML rich medium again produced a mixture of small and large colonies, the latter corresponding to spontaneous suppressor derivatives. Re-streaking of the independently isolated suppressor *ΔSGRAN_1165-sup1* (PhyR1 ΔQ194–N198) yielded uniformly large colonies, illustrating the stability of the suppressor phenotype. (c) Suppression of the *ΔSGRAN_1165* growth defect by loss of PhyR1 and/or PhyR2. Colony phenotypes of wild type TFA, *ΔSGRAN_1165ΔphyR1ΔphyR2*, *ΔSGRAN_1165ΔphyR1*, *ΔSGRAN_1165ΔphyR2*, following streaking on MML rich medium.

After several independent attempts to delete *SGRAN_1165*, two distinct colony sizes were consistently obtained following resolution of the cointegrates: large colonies corresponding to the wild type genotype and small colonies carrying the *ΔSGRAN_1165* allele (Fig. 4b). Upon re-streaking the *ΔSGRAN_1165* mutant on solid medium, a mixed population of small and large colonies was repeatedly obtained. When colonies of each morphology were independently re-streaked, large colonies maintained a stable large-colony phenotype, whereas small colonies again gave rise to a mixture of small and large colonies. The large colonies retained the *ΔSGRAN_1165* allele, indicating that they represented stable suppressor derivatives that restore normal growth.

Large colonies suppressors arising from *ΔSGRAN_1165* background isolated from independent experiments, where analyzed by whole-genome sequencing. Remarkably, all suppressor mutations mapped exclusively to the *phyR1* coding region. Three independently isolated *ΔSGRAN_1165* suppressors carried distinct alterations, including a deletion affecting residues Q194-N198 (*ΔHK_1165-sup1*), a G201V substitution (*ΔHK_1165-sup2*), and a 190 bp deletion causing frameshift from residue 123 onwards and predicted to abolish PhyR1 function, (*ΔHK_1165-sup3*). Notably, the first two alterations affect residues located close to Asp197, the predicted phospho-accepting aspartate within the REC domain of the PhyR1 response regulator, whereas the third is predicted to disrupt PhyR1 function. These findings indicate that reduction or loss of PhyR1 function alleviates the growth defect associated with the loss of HK_1165, supporting a link with the PhyR-dependent signaling.

Previous work in TFA showed that deletion of the GSR negative regulator NepR2 could not be achieved, most likely because constitutive activation of the GSR may results in a lethal overactivation of EcfG-dependent transcription (De Dios et al., 2020). Similar observations have also been reported in other alphaproteobacteria following deletion of negative regulators of the GSR (Lourenço et al., 2011; Kaczmarczyk et al., 2011; Kim et al., 2014). We therefore hypothesized that the growth defect associated with loss of HK_1165 might similarly result from excessive activation of the PhyR activity. To test this hypothesis, the *ΔSGRAN_1165* allele was introduced into a GSR-deficient background, *ΔphyR1ΔphyR2*. In contrast to the wild type, the triple mutant *ΔSGRAN_1165ΔphyR1ΔphyR2* was readily obtained and displayed colonies indistinguishable in size from those of the wild type (Fig. 4c). These results demonstrate that the deleterious phenotype associated with loss of HK_1165 depends on PhyR-mediated GSR signaling.

To determine if the observed phenotype was exclusively associated with PhyR1 or could also be achieved by eliminating PhyR2, the *ΔSGRAN_1165* allele was therefore introduced independently into *ΔphyR1* and a *ΔphyR2* backgrounds. Both *ΔSGRAN_1165ΔphyR1* and *ΔSGRAN_1165ΔphyR2* mutants were readily obtained and formed colonies of normal size (Fig. 4c). Thus, although spontaneous suppressor mutations preferentially arose in *phyR1*, genetically eliminating either PhyR paralog is sufficient to suppress the growth defect caused by loss of HK_1165. These results indicate that the deleterious phenotype of the *ΔSGRAN_1165* mutant requires the combined activity of both PhyR paralogs. These data are consistent with a model in which HK_1165 acts upstream of PhyR and establish its role as a negative regulator of PhyR-dependent GSR signaling.

### HK_1165 modulates GSR activity and stress resistance through the PhyR paralogs

To determine how HK_1165-dependent regulation of the PhyR paralogs affects GSR activity and stress adaptation, we measured GSR-dependent gene expression using the chromosomally integrated *nepR2::lacZ* reporter in the *ΔSGRAN_1165ΔphyR1* and *ΔSGRAN_1165ΔphyR2* mutants. β-galactosidase activity was measured in exponential and stationary phases and compared with that of the corresponding *ΔphyR1* and *ΔphyR2* single mutants (Fig. 5a). The three independently PhyR1 suppressor mutants were also included in the analysis (Fig. S1).

**Figure 5.**
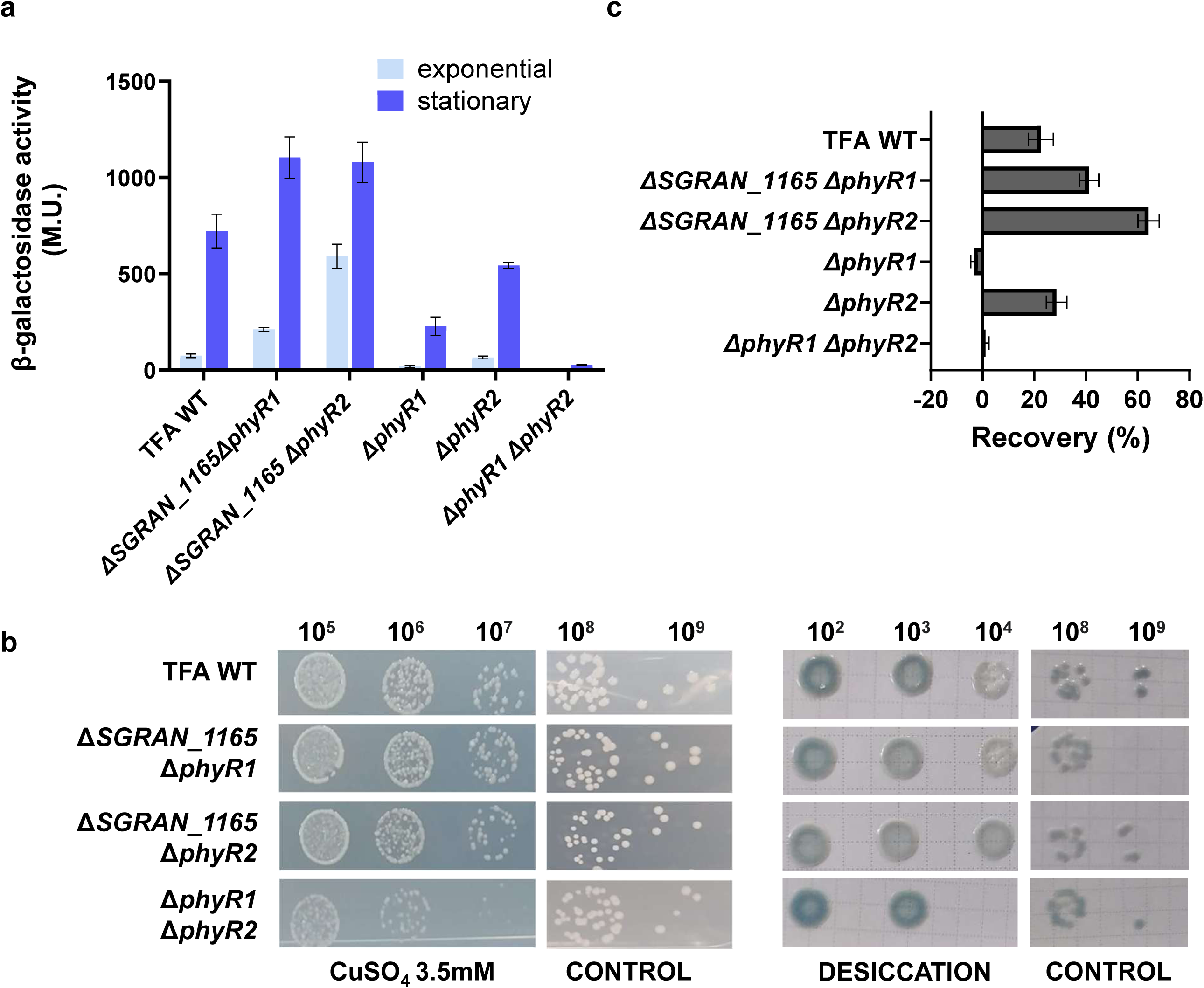
GSR activity and stress resistance phenotypes of *ΔSGRAN_1165 ΔphyR1* and *ΔSGRAN_1165 ΔphyR2* deletion mutants. (a) GSR-dependent expression was assessed during exponential growth (10 h) and stationary phase (48 h) in MM minimal medium with 8 mM β-hydroxybutyrate as the carbon source. Strains carrying a *nepR2::lacZ* translational fusion were used, and promoter activity was determined by β-galactosidase assays. (b) Heavy-metal stress sensitivity was assessed by spotting serial dilutions onto MML agar supplemented with 3.5 mM CuSO_4._ Desiccation resistance was assessed by spotting serial dilutions onto filters, air-drying them for 20 h, and subsequently placing the filters onto MML agar supplemented with 0.002% bromophenol blue. Filters transferred directly onto the plates without air-drying were used as controls. (c) Recovery from oxidative shock was assessed by comparing the OD 600 of cultures exposed to 10 mM H_2_O_2_ with that of untreated controls 3 h after treatment.

Previous work showed that the two PhyR paralogs differ in their contribution to GSR activation during stationary phase, since PhyR1 promotes stronger GSR activation than PhyR2. Accordingly, *nepR2::lacZ* expression in the *ΔphyR2* mutant, in which signaling occurs through PhyR1, was approximately threefold higher than in the *ΔphyR1* mutant, in which only PhyR2 remains functional (De Dios et al., 2022, 2020) and Fig. 5a).

Consistent with HK_1165 functioning as a negative regulator, its absence in both double mutants, *ΔSGRAN_1165ΔphyR1* and *ΔSGRAN_1165ΔphyR2*, resulted in increased GSR activation, even during exponential growth, when the pathway normally remains largely inactive. This effect was particularly evident in the *ΔSGRAN_1165ΔphyR2* mutant, which reached GSR activity levels under exponential growth comparable to the those observed in the stationary phase of wild type cells, thus activating the GSR under undesirable non stress conditions (Fig.5a). Loss of HK_1165 also markedly increased GSR activity in the *ΔphyR1* background. Whereas PhyR2 alone supported relatively low GSR activity in the *ΔphyR1* mutant (approximately 200 Miller units compared with 720 in the wild type), the *ΔSGRAN_1165ΔphyR1* mutant reached expression levels comparable to those of the wild type. Thus, in the absence of HK_1165, PhyR2 is sufficient to drive robust GSR activation, indicating that HK_1165 normally constrains PhyR2-dependent signaling. Remarkably, both *ΔSGRAN_1165ΔphyR1* and *ΔSGRAN_1165ΔphyR2*, as well as the three independent suppressor mutants, displayed resistance to copper, desiccation, and oxidative stress (Figs. 5b,c). Thus, the stress-specific contributions previously observed for single *ΔphyR* mutants were no longer evident in these backgrounds.

Whereas deletion of *phyR1* or *phyR2* alone compromises resistance to specific stresses, as illustrated by the sensitivity of the *ΔphyR1* mutant to oxidative stress and the reduced copper resistance of the *ΔphyR2* mutant, elimination of HK_1165 restored broad stress resistance even when only a single PhyR paralog remained functional. This indicates that, in the absence of HK_1165, either PhyR paralog alone is sufficient to sustain resistance to all three stresses tested. These results further suggest that the apparent functional specialization of PhyR1 and PhyR2 depends not only on its intrinsic properties but also on their phosphorylation state. In the absence of HK_1165, elevated phosphorylation levels of the remaining PhyR paralog may override this specialization, allowing PhyR protein to drive a more generalized GSR output.

Taken together these expression data and phenotypic analyses, indicate that HK_1165 functions as a major negative regulator of the GSR that impedes overactivation of the system potentially acting as a PhyR phosphatase. In addition to prevents inappropriate GSR activation under non-stress conditions, HK_1165 modulates the activity of each PhyR paralog, determining their relative contribution to different stress responses.

### HK1165 controls GSR output by modulating PhyR phosphorylation *in vivo*

Based on the increased GSR output and stress resistance observed in the absence of HK_1165, we next investigated whether these phenotypes correlate with changes in the phosphorylation state of the PhyR paralogs *in vivo*.

To this end, the phosphorylation state of PhyR1 and PhyR2 was assessed *in vivo* using Phos-tag SDS-PAGE, which enables the separation of phosphorylated and unphosphorylated protein forms based on mobility shifts. For specific detection, chromosomally 3XFLAG-tagged versions of PhyR1 and PhyR2 were constructed in the corresponding genetic backgrounds and detected by immunoblotting. PhyR phosphorylation was analysed in the WT background and, to specifically assess PhyR2 phosphorylation and the effect of HK_1165 on this paralog, in the *ΔphyR1* and *ΔSGRAN_1165ΔphyR1* backgrounds. PhyR phosphorylation was examined by Western blot under oxidative stress conditions following H_2_O_2_ exposure, a condition in which previous genetic analyses showed a predominant contribution of PhyR1 to GSR signaling and upon stationary phase an GSR inducing condition in TFA (De Dios et al., 2022).

In the WT strain, exposure to H_2_O_2_ increased the proportion of phosphorylated PhyR1, which was the predominant phosphorylated species, consistent with the major contribution of this paralog to GSR signalling under oxidative stress (Fig. 6a,b). PhyR2 remained largely unphosphorylated in the *ΔphyR1* mutant, suggesting that this paralog contributes only minimally to GSR signalling under oxidative stress.

**Figure 6.**
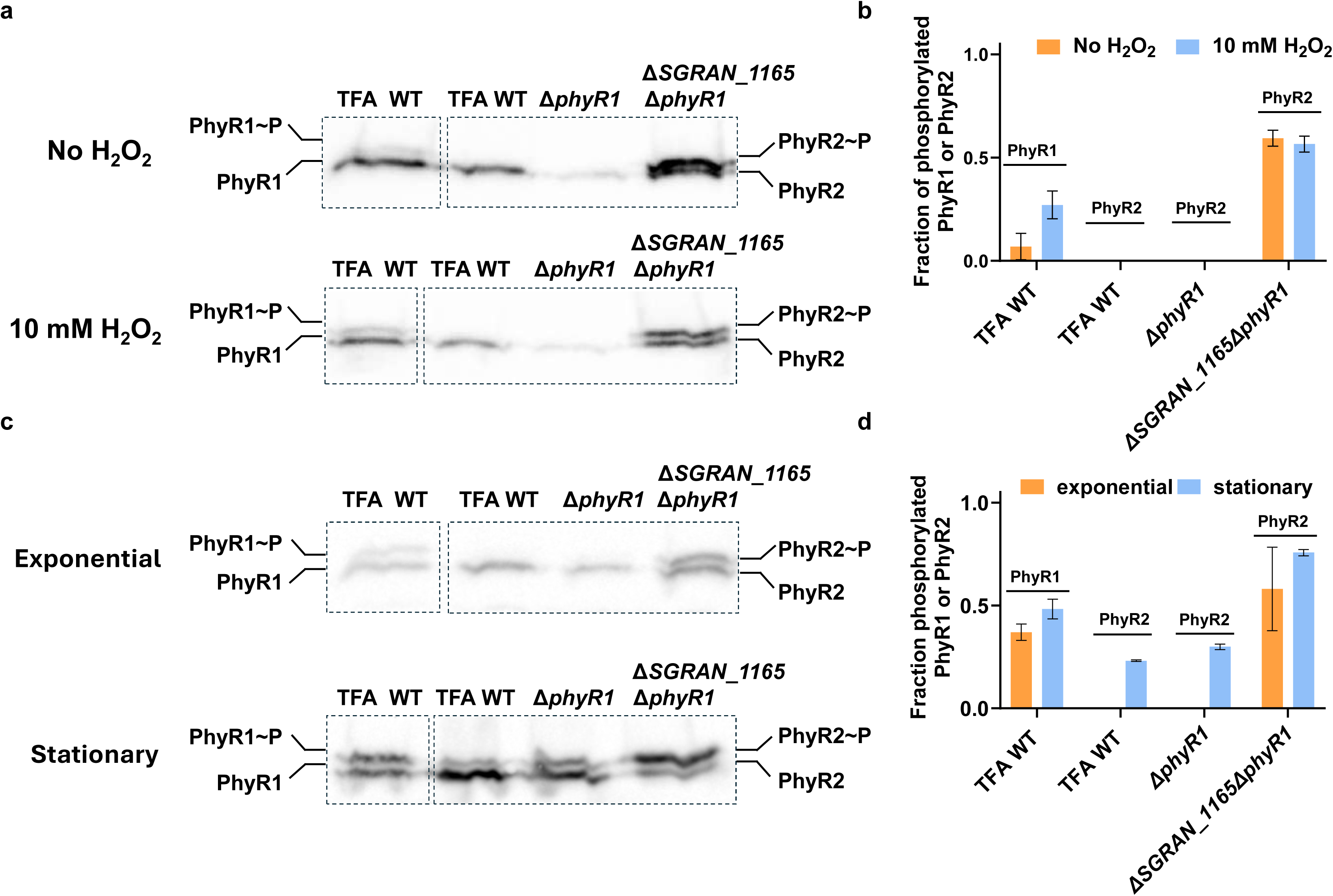
Analysis of PhyR1 and PhyR2 phosphorylation *in vivo* by Phos-tag SDS-PAGE. Western blot detection of PhyR1 and PhyR2 protein under different conditions in TFA WT, *ΔphyR1* and *ΔSGRAN_1165 ΔphyR1* strains after electrophoresis in a 25µM phos-tag tm acrylamide gel. (a) Representative Western blot analysis of PhyR1 and PhyR2 phosphorylation in wild-type TFA, *ΔphyR1*, and *ΔSGRAN_1165ΔphyR1* strains before and after exposure to 10 mM H_2_O_2_ for 3 h. Proteins were separated by SDS-PAGE using gels containing 25 µM Phos-tag acrylamide and detected by immunoblotting with anti-FLAG antibodies. Phosphorylated PhyR1 and PhyR2 migrated more slowly than their corresponding unphosphorylated forms. (b) two independent Western blot were quantified and the fraction of phosphorylated PhyR was calculated relative to the total signal of phosphorylated and unphosphorylated forms of each protein. Labels above the bars indicate the PhyR paralog quantified in each strain. When no phosphorylated band was detected, the fraction was assigned a value of 0. (c) Representative Western blot analysis of PhyR1 and PhyR2 phosphorylation in exponential and stationary phases in cultures grown in minimal medium with 8 mM β-hydroxybutyrate (BHB) as carbon source. (d) Quantification of PhyR phosphorylation from two independent Western blot experiments performed as described in (b).

However, deletion of HK_1165 in the *ΔphyR1* background increased the proportion of phosphorylated PhyR2, both in the absence and presence of H_2_O_2_ (Fig. 6a,b). This increase was consistent with the enhanced GSR activity and recovery of oxidative stress resistance observed in the *ΔSGRAN_1165ΔphyR1* mutant (Fig. 5) These results suggests that, when HK_1165 is absent, increased accumulation of PhyR2∼P enables PhyR2 alone to sustain robust GSR activation and oxidative stress resistance. Thus, the limited GSR activation supported by PhyR2 under these conditions appears to be associated with low levels of its phosphorylated form rather than simply with a lower intrinsic capacity of PhyR2 to activate the GSR.

A similar pattern was observed during stationary phase, where loss of HK_1165 in the *ΔphyR1* background also promoted marked accumulation of phosphorylated PhyR2. This increase closely correlates with having enhanced GSR-dependent *nepR2::lacZ* expression observed in the *ΔSGRAN_1165ΔphyR1* mutant, compared with the *ΔphyR1* single mutant (approximately 5-fold higher, Fig. 5a).

These findings provide a mechanistic basis for the phenotypic and reporter analyses presented above. HK_1165 preferentially limits PhyR2 phosphorylation under oxidative stress and stationary phase conditions, thereby restricting PhyR2-dependent GSR activation and favouring signalling through PhyR1. When HK_1165 is absent, PhyR2 accumulates in its phosphorylated form and becomes sufficient to support GSR activation under conditions in which this paralog normally contributes only weakly to signalling. This increased PhyR2 activity is consistent with the ability of the *ΔHK_1165ΔphyR1* mutant to withstand stresses, including oxidative stress and even desiccation, for which PhyR2 alone normally provides limited protection. This fact also would explain why PhyR2 alone can support resistance not only to oxidative stress but also to desiccation, a condition that normally requires the coordinated contribution of both PhyR paralogs.

More generally, these data support a model in which HK_1165 controls the amount of phosphorylated PhyR available for GSR activation, thereby influencing the relative contribution of each PhyR paralog to the stress response.

### HK_3483 phosphorylates both PhyR paralogs with a preference for PhyR1

The genetic analyses described above identified HK3483 as a major positive regulator of the GSR involved in sensing multiple environmental stresses mediated by the two PhyR response regulators. These pointed out a role of HK_3483 to promote phosphorylation of both response regulators at their REC domains.

To validate this function, we performed *in vitro* phosphorylation assays using purified HK_3483, PhyR1 and PhyR2 proteins and radiolabeled ATP as indicated in Material and Methods.

Since HK_3483 is a membrane-associated histidine kinase containing an N-terminal CHASE sensory domain flanked by transmembrane helices, we generated a truncated variant lacking the CHASE domain and its flanking transmembrane helices (residues 1-337) to facilitate the production of a soluble protein for biochemical analysis. Similar truncations have previously been shown to retain autokinase and phosphotransfer activities in other CHASE-containing histidine kinases (Kim et al., 2014; Najnin et al., 2016). HK_3483 truncated of a predicted molecular mass of approximately 24kDa was expressed and purified in soluble form. Purified protein (10µM) was used to assess its autokinase activity using [γ-32P] ATP as a phosphodonor. A time-course analysis showed progressive accumulation of phosphorylated HK_3483, reaching a steady state approximately 15 min after initiation of the reaction (Fig. 7a).

**Figure 7:**
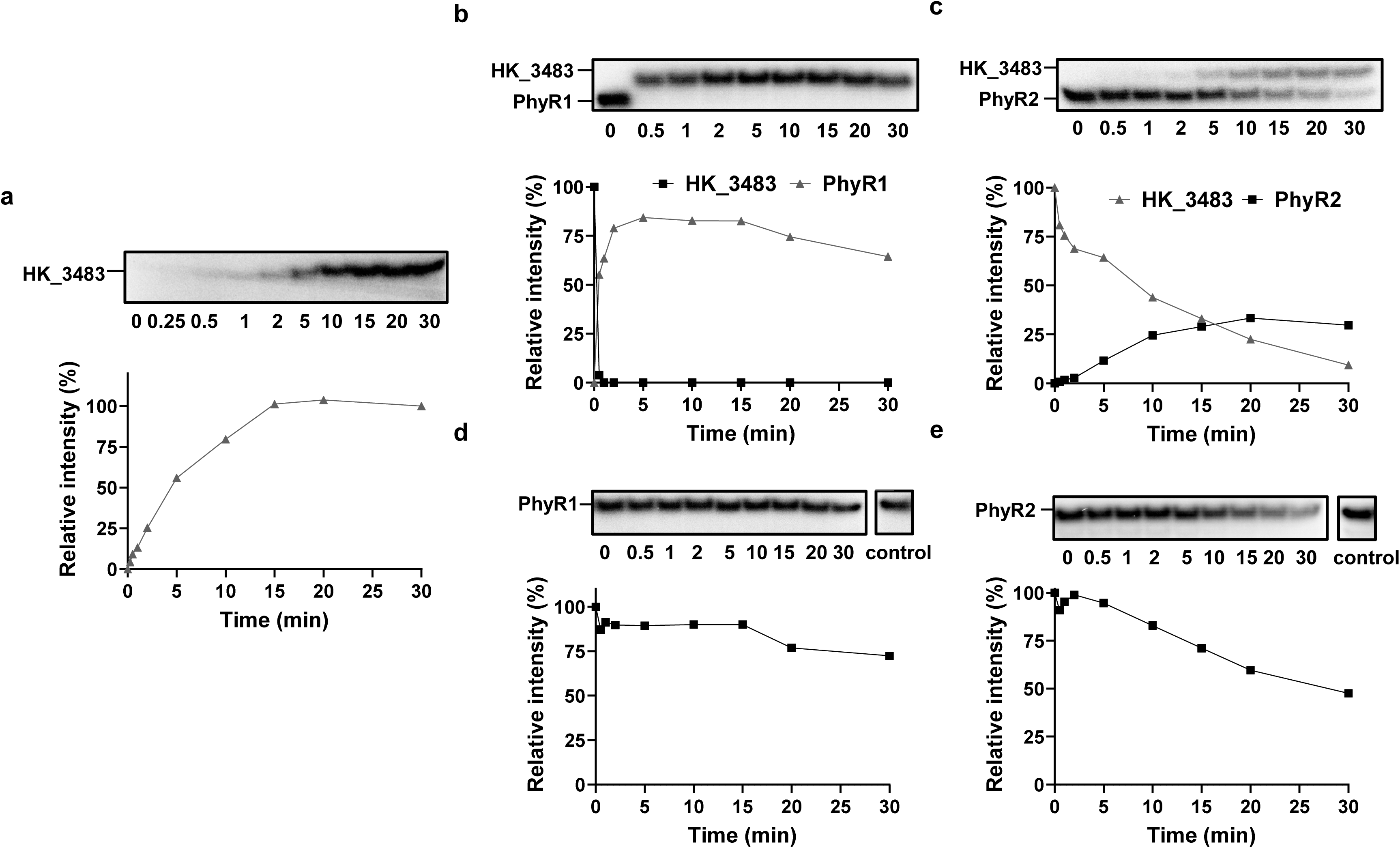
*In vitro* phosphorylation and dephosphorylation assays of PhyR1 and PhyR2 by either HK_3483 or HK1165 respectively. (a) Autokinase activity of the truncated HK_3483 protein in the presence of [γ-32P] ATP. HK_3483∼P signal shows saturation after 15 minutes of incubation. The HK_3483∼P signal at 30 min was set to 100% for quantification. Phosphotransfer from pre-phosphorylated HK_3483 to PhyR1 (b) or PhyR2 (c). Phosphotransfer to PhyR1 occurs rapidly with an almost complete loss of the HK_3483∼P signal within 30 s, whereas phosphotransfer to PhyR2 proceeded considerably more slowly. Signals corresponding to HK_3483∼P and PhyR∼P were expressed relative to the HK_3483∼P signal at 0 min, which was set to 100%. Quantification of HK_3483 autophosphorylation and phosphotransfer is shown alongside the corresponding autoradiograms. (d, e) Dephosphorylation of pre-phosphorylated PhyR1 (d) or PhyR2 (e) in the presence of full-length HK_1165. PhyR1∼P levels remained largely unchanged throughout the assay and were comparable to those in the control reaction lacking HK_1165. In contrast, PhyR2∼P progressively decreased in the presence of HK_1165 compared with the control reaction. For quantification, the PhyR1∼P or PhyR2∼P signal at 0 min was set to 100%.

Next, to determine whether HK_3483 directly phosphorylates the two PhyR paralogs, phosphotransfer reactions were carried out using equimolar concentrations (5µM) of autophosphorylated HK_3483 and purified PhyR1 or PhyR2 (Fig. 7b,c). Phosphotransfer to PhyR1 occurs very rapidly, with a complete loss of HK_3483 phosphorylation signal within 30 seconds after mixing the proteins, concomitant with the rapid accumulation of phosphorylated PhyR1 (Fig. 7b). In contrast, phosphotransfer to PhyR2 proceeded more slowly. Phosphorylated HK_3483 decreased progressively over the course of the reaction, concomitant with the accumulation of phosphorylated PhyR2, with most of the HK_3483 phosphorylation signal lost after 30 min (Fig. 7c). Thus, HK_3483 directly phosphorylated both PhyR paralogs, but with different phosphotransfer kinetics.

These results demonstrate that HK_3483 is capable of autophosphorylation and functions as a phosphodonor for both PhyR paralogs, indicating that HK_3483 can directly signal to PhyR1 and PhyR2, providing a biochemical basis for the increased sensitivity of the *ΔSGRAN_3483* mutant to the environmental stresses tested.

### HK_1165 exhibits phosphatase activity primarily toward PhyR2 *in vitro*

The difficulty in obtaining an HK_1165 deletion unless PhyR-dependent GSR signalling was impaired suggest that a tight control of PhyR activity by HK_1165 is important to prevent excessive GSR activation. Also, consistent with this idea, loss of HK_1165 increased PhyR phosphorylation *in vivo*. Together, these observations prompted us to investigate whether HK_1165 could act as a phosphatase toward the two PhyR paralogs.

HK_1165 was overproduced in *E. coli* and purified as a soluble recombinant protein of approximately 57kDa.To assess the phosphatase activity, PhyR1 and PhyR2, were first phosphorylated *in vitro* using HK_3483 and [γ-32P] ATP and subsequently incubated with purified HK_1165. The decay of the radiolabeled PhyR∼P signal was then monitored over time.

When PhyR1∼P was incubated with HK_1165 no appreciable decrease in phosphorylation signal was observed over the 30 min of incubation period, (Fig. 7d), whereas in the case of PhyR2∼P, a progressive decrease in PhyR2∼P became apparent after approximately 10 min of incubation with HK_1165 (Fig. 7e). No concomitant accumulation of phosphorylated HK_1165 was detected during the reaction. These results indicate that HK_1165 exhibits phosphatase activity *in vitro*, with a strong preference for PhyR2.

Although the *in vitro* assays reveal a clear preference of HK_1165 for PhyR2, our *in vivo* analysis indicate that HK_1165 influences signalling through both PhyR paralogs. In particular, the elevated GSR-dependent *nepR2::lacZ* expression observed in the *ΔSGRAN_1165ΔphyR2* mutant (Fig. 5a) during exponential growth indicates that HK_1165 also restricts PhyR1-dependent GSR activation *in vivo*.

Together with the phosphotransfer assays showing that HK_3483 phosphorylates PhyR1 more rapidly than PhyR2, these results reveal a functional asymmetry in the system whereby HK_3483 preferentially drives PhyR1 phosphorylation while HK_1165 preferentially removes phosphate from PhyR2. These opposing regulatory preferences likely contribute to establishing distinct phosphorylation balances between the two PhyR paralogs, thereby shaping their relative contribution to GSR activation under different physiological conditions.

## Discussion

The general stress response (GSR) in alphaproteobacteria relies on signalling networks that control the phosphorylation state of the response regulator PhyR, thereby modulating the activity of EcfG sigma factors. Although several HWE/HisKA2-family histidine kinases have been implicated in GSR regulation (Kaczmarczyk et al., 2014; Kim et al., 2014; Lori et al., 2018), the mechanisms coordinating pathway activation and attenuation remain incompletely understood, particularly in organisms possessing multiple PhyR paralogs.

*Sphingopyxis granuli* TFA is a pollutant-degrading alphaproteobacterium in which stress resistance is controlled by multiple regulatory pathways. Our recent work showed that the small RNA SuhB contributes to stress resistance through mechanisms largely independent of the GSR (García-Romero et al., 2026).

Within the GSR, TFA possesses two PhyR paralogs with distinct contributions to GSR activation and stress resistance (De Dios et al., 2022, 2020). Previous work showed that PhyR1 makes a greater contribution than PhyR2 to GSR-dependent gene expression during stationary phase, as revealed by the different reporter activities of the corresponding single-deletion mutants. The relative contributions of the two paralogs also vary with the stress:: PhyR1 is particularly important for resistance to oxidative stress, whereas PhyR2 can sustain copper resistance in the absence of PhyR1, and both paralogs contribute to full desiccation resistance (De Dios et al., 2022). These observations raise the question of how the activities of the two PhyR paralogs are differentially regulated and whether this regulation contributes to their functional specialization.

Here, we identified two HWE/HisKA2-family proteins with distinct domain architectures, the membrane-associated with a CHASE-domain histidine kinase HK_3483 and the predicted cytoplasmic GAF-PAS-containing protein HK_1165, as major positive and negative regulators of the GSR, respectively. Genetic and biochemical analyses established that HK_3483 is required for effective GSR activation through both PhyR paralogs, whereas HK_1165 restricts PhyR-dependent signalling and prevents excessive pathway activation. Our findings suggest that the opposing activities of HK_3483 and HK_1165 contribute to establishing the phosphorylation balance between the two PhyR paralogs, thereby determining their relative contributions to GSR activation and stress resistance.

### HK_3483 is a major upstream activator of both PhyR paralogs

The association between HWE/HisKA2-family kinases and PhyR-dependent GSR signalling is supported by their distribution across Alphaproteobacteria. The reported absence of HRXXN-containing kinases in species lacking the PhyR–NepR–EcfG cascade suggests an evolutionary link between these signalling components (Kaczmarczyk et al., 2014). Although the core partner-switching mechanism is conserved, GSR regulatory networks differ in the number and functional organization of their components. Additional EcfG sigma factors can operate hierarchically, as in *Caulobacter crescentus* (Lourenço et al., 2011) or in parallel, as in *Methylobacterium extorquens* (Francez-Charlot et al., 2016), whereas additional NepR proteins can introduce negative feedback, as demonstrated in *Sphingomonas melonis* Fr1 (Gottschlich et al., 2019). *In S. granuli TFA*, this regulatory diversification extends to all three core components, with two EcfG sigma factors, two NepR anti-sigma factors and two PhyR response regulators (De Dios et al., 2022, 2020). Unlike the functional redundancy reported for the two PhyR-like regulators of *Sinorhizobium meliloti* (Bastiat et al., 2010) PhyR1 and PhyR2 in TFA make distinct contributions to stress resistance (De Dios et al., 2022). This raises the question of how upstream regulators control the activity of each paralog and contribute to their functional specialization.

Among the four candidate HWE/HisKA2 histidine kinases identified in TFA, HK_3483 emerged as the major positive regulator of the GSR under the conditions examined. Several complementary findings support this role. Loss of HK_3483 compromises resistance to copper, oxidative stress and desiccation and markedly reduces GSR-dependent reporter activity (Fig.2). Genetic analysis of mutants lacking HK_3483 and either PhyR paralog indicate that efficient signalling through the remaining paralog requires HK_3483 (Fig. 3). Together with the direct phosphorylation of both PhyR proteins by HK_3483 in vitro (Fig.7), establish HK_3483 as a shared upstream phosphodonor for the two response regulators.

The contribution of HK_3483 to resistance to three distinct stresses suggests that this sensor histidine kinase participates in transmitting signals associated with different environmental challenges. This is consistent with the overlapping contributions of individual PhyR-activating kinases described in *S. melonis* Fr1, where a single kinase can participate in responses to more than one stress (Kaczmarczyk et al., 2014). However, whereas partial redundancy distributes GSR activation among several Paks in that organism, the pronounced defects caused by loss of HK_3483 indicate a predominant role for this kinase under the conditions examined in TFA. Whether these stresses influence HK_3483 through a common cellular signal or through distinct mechanisms remains to be determined. Nevertheless, its preference for PhyR1 *in vitro* suggests that its shared role does not imply equivalent input to both paralogs.

### HK_1165 negatively regulates GSR signalling and prevents excessive pathway activation

In contrast to HK_3483, HK_1165 functions as a major negative regulator of the GSR. The pronounced growth defect associated with deletion of HK_1165, together with the emergence of spontaneous suppressor mutants, suggests that maintaining appropriate levels of GSR activity is important for cellular fitness. A comparable requirement for negative regulation has been documented in S. melonis Fr1, where PhyT, formerly designated PhyP, prevents lethal GSR overactivation. Deletion of this gene could be obtained in backgrounds with impaired GSR signalling, such as mutants lacking PhyR or EcfG regulators (Gottschlich et al., 2018; Kaczmarczyk et al., 2011). The duplicated PhyR system of TFA provides an opportunity to examine this requirement of negative regulation while still retaining PhyR-dependent signalling. Deletion of either phyR1 or phyR2 suppresses the growth defect associated with loss of HK_1165, even though the remaining paralog supports elevated GSR activity and broad stress resistance (Figs. 4 and 5). Thus, suppression does not require complete inactivation of PhyR-dependent signalling. Instead, deleterious effect of HK_1165 loss depends on the combined activity of both PhyR paralogs. The recovery of three independent suppressors carrying mutations in the *phyR1* further links the growth defect to deregulated PhyR activation (Fig. 4). Together, these results suggest that HK_1165 constrains the combined output of the two PhyR branches, allowing protective GSR activation while preventing signalling levels that compromise growth. Negative regulators of the GSR have been identified in several alphaproteobacteria, but their mechanisms of action are not always fully resolved. In *S. melonis* Fr1, PhyT, functions as a phosphotransferase in the GSR-activating Pak–SdrG–PhyT–PhyR phosphorelay. Although initially proposed to act as a PhyR phosphatase (Kaczmarczyk et al., 2011), its negative regulatory function was subsequently attributed to complex formation with unphosphorylated PhyR, which restricts inappropriate phosphorylation. Direct phosphatase activity towards phosphorylated PhyR has not been demonstrated (Gottschlich et al., 2018). Similarly, the HWE-family protein RsiC contributes to both positive and negative regulation of the GSR in *S. meliloti*. Its proposed phosphatase function was supported by genetic and *in vivo* phosphorylation analyses but could not be tested biochemically (Sauviac and Bruand, 2014).

Our results provide a biochemical mechanism by which HK_1165 can restrict PhyR activation since HK_1165 clearly promotes the loss of phosphate from PhyR2∼P *in vitro* (Fig.7). Under the same experimental conditions, PhyR1∼P remains largely unaffected, suggesting that negative regulation can act differentially on the two paralogs.

### Differential control of PhyR phosphorylation shapes the contributions of the two paralogs to GSR signalling

The opposing biochemical activities of HK_3483 and HK_1165 are consistent with the phosphorylation patterns observed *in vivo* under oxidative stress and during stationary phase, providing a possible mechanistic explanation for the predominant involvement of PhyR1 under these stress signalling. HK_3483 preferentially phosphorylates PhyR1, whereas HK_1165 preferentially promotes PhyR2 dephosphorylation *in vitro*. Consistent with these activities, Phos-tag analyses revealed a higher proportion of phosphorylated PhyR1 than PhyR2 in wild-type cells under oxidative stress and during stationary phase (Fig.6). Moreover, deletion of HK_1165 in the *ΔphyR1* background resulted in increased accumulation of PhyR2∼P, resulting in restored H_2_O_2_ resistance and enhanced GSR activity (Figs 5 and 6), Thus, the limited contribution of PhyR2 to oxidative stress resistance under normal regulatory conditions, cannot be explained by an intrinsic inability to support this response; instead, it reflects, at least in part, its restricted phosphorylation. These observations support a model in which the opposing activities of HK_3483 and HK_1165 contribute to establishing an phosphorylation pattern between the two PhyR paralogs, favouring signalling through PhyR1 under the conditions examined.

Although HK_1165 preferentially dephosphorylates PhyR2∼P *in vitro*, our genetic results indicate that *in vivo* its negative regulatory influence could also extend to PhyR1. In the *ΔHK_1165ΔphyR2* mutant, where PhyR1 is the only remaining PhyR protein, GSR reporter activity was markedly increased, reaching levels during exponential growth (non-stress conditions) comparable to those observed in stationary phase (Fig 5). The appearing of independent suppressor mutations in PhyR1 further supports the importance of controlling PhyR1 activity in the absence of HK_1165. This indicate that HK_1165 may also promote PhyR1 dephosphorylation *in vivo*, contributing to the growth-phase-dependent regulation of GSR despite the absence of detectable biochemical activity towards PhyR1∼P. This could reflect a condition not reproduced in our *in vitro* assays, or it could be an indirect consequence of altered GSR regulation.

The consequences of HK_1165 deletion extend beyond oxidative stress resistance or stationary phase. Under normal regulatory conditions, PhyR2 is particularly important for copper resistance, whereas full desiccation resistance requires contributions from both paralogs (De Dios et al., 2022). Both *ΔHK_1165ΔphyR1* and *ΔHK_1165ΔphyR2* exhibit resistance to copper, oxidative stress and desiccation, despite retaining only one PhyR paralog (Fig. 5). Thus, the stress-specific limitations observed in the corresponding single *ΔphyR* mutants are no longer apparent when HK_1165 is absent, indicating that either paralog can sustain broad stress resistance under altered regulatory conditions. Although PhyR phosphorylation has not been examined under copper or desiccation stress, these phenotypes suggest that HK_1165-dependent regulation also influences the contributions of the two paralogs to protection against these stresses.

Diversification of sensory histidine kinases is a way to expand the range of signals integrated by alphaproteobacterial GSR networks while maintaining convergence on a common PhyR regulator (Kaczmarczyk et al., 2014). The plasticity and modular organization of these upstream networks may facilitate adaptation to different ecological niches (Lori et al., 2018). Our findings highlight an additional dimension of this regulatory organization: a shared upstream network can differentially control two PhyR paralogs contributing to their functional specialization. Thus, diversification of sensory inputs and differential regulation of response regulator phosphorylation may represent complementary means of shaping GSR signalling.

Together, these findings support a model in which HK_3483 and HK_1165 act as opposing regulatory components of the GSR signalling network in TFA. Their differential control of PhyR phosphorylation not only helps determine whether the pathway is active or inactive, but also shapes the phosphorylation state of each PhyR paralog and, thereby, its contribution to GSR activation and stress resistance, providing a regulatory basis for their functional specialization.

## Acknowledgements

We are grateful to the members of the Division of Microbiology at the University Pablo de Olavide (Seville, Spain) for their advice and suggestions, and to the laboratory technicians for their invaluable technical support. We also thank the proteomics facility at CABD for their assistance.

