## Supplementary_materials for "An HWE/HisKA2-family kinase-phosphatase pair differentially regulates two paralogous PhyR response regulators to shape the General Stress Response specificity in *Sphingopyxis granuli* TFA"

Centro Andaluz de Biología del Desarrollo, Universidad Pablo de Olavide/Consejo Superior de Investigaciones Científicas/Junta de Andalucía, ES-41013 Seville, Spain

### **Supplementary Table S1**: Antibiotic concentration for *E. coli* and *S. granuli* TFA

| **Species** | **Antibiotic** | **Concentration (µg/mL)** |
|---|---|---|
| ***E. coli*** | Kanamycin | 25 |
| | Ampicillin | 100 |
| ***S. granuli* TFA** | Kanamycin | 20 |
| | Ampicillin | 5 |
| | Streptomycin | 50 or 200 |

### **Supplementary Table S2**: Strains and plasmids used in this study.

| **Strains** | **Description** | **Reference** |
|---|---|---|
| ***Escherichia coli*** | | |
| **DH5α** | F- *ø80lacZΔM15 Δ(lacZYA-argF) U169, recA1, endA1 hsdR17(rK-mK$^-$), supE44, thi-1, gyrA, relA1* | (Hanahan, 1983) |
| **DH5α λ*pir*** | λ*pir* phage lysogen of DH5α | Lab collection |
| **ER2566** | F- λ- *fhuA2, [lon], ompT, lacZ::T7p07, gal, sulA11,* Δ*(mcrCmrr)114::IS10,* R*(mcr-73::miniTn10-TetS)2,* R*(zgb210::Tn10)(TetS), endA1, [dcm]* | New England Biolabs |
| ***Sphingopyxis granuli* TFA** | | |
| **TFA WT** | Wild type. Str$^r$. | (Hernáez et al., 1999) |
| **MPO858** | *nepR2::lacZ* translational fusion integrated into TFA WT chromosome. Ap$^r$, Str$^r$ | (De Dios et al., 2020) |
| **MPO860** | *ecfG1* and *ecfG2* double deletion mutant. Str$^r$ | (De Dios et al., 2020) |

| | | |
|---|---|---|
| **MPO866** | *phyR1* deletion mutant. $Str^r$ | (De Dios et al., 2022) |
| **MPO867** | *phyR2* deletion mutant. $Str^r$ | (De Dios et al., 2022) |
| **MPO868** | *phyR1* and *phyR2* double deletion mutant. $Str^r$ | (De Dios et al., 2022) |
| **MPO872** | *nepR2::lacZ* translational fusion integrated into MPO866 chromosome. $Ap^r$, $Str^r$ | This work |
| **MPO873** | *nepR2::lacZ* translational fusion integrated into MPO867 chromosome. $Ap^r$, $Str^r$ | This work |
| **MPO874** | *nepR2::lacZ* translational fusion integrated into MPO868 chromosome. $Ap^r$, $Str^r$ | This work |
| **MPO910** | *phyR1::3xflag* translational fusion integrated into TFA WT chromosome. $Str^r$ | (De Dios et al., 2022) |
| **MPO911** | *phyR2::3xflag* translational fusion integrated into TFA WT chromosome. $Str^r$ | (De Dios et al., 2022) |
| **MPO924** | *SGRAN_2544* deletion mutant. $Str^r$. | This work |
| **MPO925** | *nepR2::lacZ* translational fusion integrated into MPO924 chromosome. $Ap^r$, $Str^r$ | This work |
| **MPO927** | *SGRAN_3483* deletion mutant. $Str^r$ | This work |
| **MPO928** | *SGRAN_1773 deletion mutant.* $Str^r$ | This work |
| **MPO929** | *SGRAN_1773* and SGRAN 3483 double deletion mutant. $Str^r$ | This work |
| **MPO930** | *SGRAN_1165, phyR1* and *phyR2* triple deletion mutant. $Str^r$ | This work |
| **MPO931** | *SGRAN_2544* and *SGRAN 3483* double deletion mutant. $Str^r$ | This work |
| **MPO932** | *SGRAN_1773* and *SGRAN 2544* double deletion mutant. $Str^r$ | This work |
| **MPO933** | *SGRAN_1165* and *phyR1* double deletion mutant. $Str^r$ | This work |
| **MPO934** | *SGRAN_1165* and *phyR2* double deletion mutant. $Str^r$ | This work |
| **MPO936** | SGRAN_1773, *SGRAN_2544* and *SGRAN 3483* triple deletion mutant. $Str^r$ | This work |
| **MPO937** | *SGRAN_3483* and *phyR1* double deletion mutant. $Str^r$ | This work |
| **MPO938** | *SGRAN_3483* and *phyR2* double deletion mutant. $Str^r$ | This work |
| **MPO941** | *phyR2::3xflag* translational fusion integrated into MPO866 chromosome. $Str^r$ | This work |
| **MPO943** | *phyR2::3xflag* translational fusion integrated into MPO933 chromosome. $Str^r$ | This work |
| **MPO945** | *nepR2::lacZ* translational fusion integrated into MPO928 chromosome. $Ap^r$, $Str^r$ | This work |
| **MPO947** | *nepR2::lacZ* translational fusion integrated into MPO927 chromosome. $Ap^r$, $Str^r$ | This work |
| **MPO948** | *nepR2::lacZ* translational fusion integrated into MPO932 chromosome. $Ap^r$, $Str^r$ | This work |
| **MPO949** | *nepR2::lacZ* translational fusion integrated into MPO929 chromosome. $Ap^r$, $Str^r$ | This work |
| **MPO1001** | *nepR2::lacZ* translational fusion integrated into MPO931 chromosome. $Ap^r$, $Str^r$ | This work |
| **MPO1002** | *nepR2::lacZ* translational fusion integrated into MPO936 chromosome. $Ap^r$, $Str^r$ | This work |

| | | |
|---|---|---|
| **MPO1003** | *nepR2::lacZ* translational fusion integrated into MPO937 chromosome. $Ap^r$, $Str^r$ | This work |
| **MPO1004** | *nepR2::lacZ* translational fusion integrated into MPO938 chromosome. $Ap^r$, $Str^r$ | This work |
| **MPO1005** | *nepR2::lacZ* translational fusion integrated into MPO933 chromosome. | This work |
| **MPO1006** | *nepR2::lacZ* translational fusion integrated into MPO934 chromosome. $Ap^r$, $Str^r$ | This work |
| **MPO1010** | Δ*SGRAN_1165* - sup1. It contains a deletion affecting residues Q194-N198 of the *phyR1* gene. $Str^r$ | This work |
| **MPO1011** | Δ*SGRAN_1165* – sup2. It contains a G201V substitution affecting the *phyR1* gene. $Str^r$ | This work |
| **MPO1012** | Δ*SGRAN_1165* – sup3. It contains a frame-shifting deletion of 190 nucleotides in the *phyR1* gene. $Str^r$ | This work |
| **Plasmids** | **Description** | **Reference** |
| **pMPO1408** | pJES379 derivative bearing a *nepR2::lacZ* translational fusion. $Ap^r$ | (De Dios et al., 2020) |
| **pMPO1412** | pEMG derivative with *rpsL1* streptomycin counterselection marker. $Km^r$. | (García-Romero et al., 2024) |
| **pMPO1436** | pTXB1 derivative bearing intein::*phyR1* fusion for PhyR1 purification. $Ap^r$ | (De Dios et al., 2022) |
| **pMPO1437** | pTXB1 derivative bearing intein::*phyR2* fusion for PhyR2 purification. $Ap^r$ | (De Dios et al., 2022) |
| **pMPO1459** | pMPO1412 derivative bearing flanking region C-terminal 3×FLAG tagging of PhyR1. $Km^r$ | (De Dios et al., 2022) |
| **pMPO1460** | pMPO1412 derivative bearing flanking regions for C-terminal 3×FLAG tagging of PhyR2. $Km^r$ | (De Dios et al., 2022) |
| **pMPO1469** | pMPO1412 derivative bearing *SGRAN_1165* flanking regions for its deletion. $Km^r$ | This work |
| **pMPO1470** | pMPO1412 derivative bearing *SGRAN_2544* flanking regions for its deletion, $Km^r$ | This work |
| **pMPO1471** | pMPO1412 derivative bearing *SGRAN_1773* flanking regions for its deletion, $Km^r$ | This work |
| **pMPO1473** | pTYB21 derivative bearing intein:: truncated *SGRAN_3483* fusion for protein purification. $Ap^r$. | This work |
| **pMPO1474** | pTYB21 derivative bearing intein::*SGRAN_1165* fusion for HK_1165 purification. $Ap^r$ | This work |
| **pMPO1480** | pMPO1412 derivative bearing SGRAN_3483 flanking regions for its deletion, $Km^r$ | This work |
| **pSW-I** | oriRK2, *xylS*, Pm::*I-SceI*. $Ap^r$ | (Martínez-García and de Lorenzo, 2011) |
| **pTYB21** | *E. coli* expression vector containing an N-terminal intein fusion for protein purification using the IMPACT system kit, $Ap^r$. | New England Biolabs |

**Supplementary Table S3**: Oligonucleotides used in this study.

| Oligonucleotide | Sequence | Use |
|---|---|---|
| **Sgran2544_1** | atattctagaAGGAGAGCCAGGATCAGG | PCR of upstream region of *SGRAN_2544* |
| **Sgran2544_2** | atatggatccCATAAGCGCGCCCAGGC | PCR of upstream region of *SGRAN_2544* |
| **Sgran2544_3** | tataggatccGTTCCAAAATCGCGGGAG | PCR of downstream region of *SGRAN_2544* |
| **Sgran2544_5** | tatacccgggcTTCACGGACATGAGTTCG | PCR of downstream region of *SGRAN_2544* |
| **Sgran1773_1** | atattctagaCGCCGCTCGTCAACGTCA | PCR of upstream region of *SGRAN_1773* |
| **Sgran1773_2** | atatggatccCCGGCGCATGCCATCGTC | PCR of upstream region of *SGRAN_1773* |
| **Sgran1773_3** | tataggatccTCCCACGAGCTTAAGGTG | PCR of downstream region of *SGRAN_1773* |
| **Sgran1773_4** | tatacccgggcATGAGAATTCCCGTCGTG | PCR of downstream region of *SGRAN_1773* |
| **Sgran3483_1** | ATATTCTAGAATATCGACGATTTCTGGG | PCR of upstream region of *SGRAN_3483* |
| **Sgran3483_2** | ATATGGATCCCATTGCTTGGTTTTCCTG | PCR of upstream region of *SGRAN_3483* |
| **Sgran3483_5** | TATAGGATCCCTTGCCATCCTGATCCGC | PCR of downstream region of *SGRAN_3483* |
| **Sgran3483_6** | TATACCCGGGCGTGTGGATGCGCAGCATC | PCR of downstream region of *SGRAN_3483* |

| | | |
|---|---|---|
| **Sgran1165_1** | ATATTCTAGAGTTCTATAGCCTGTGGGG | PCR of upstream region of *SGRAN_1165* |
| **Sgran1165_2** | ATATGGATCCCCGCTTTGCCATGAATCC | PCR of upstream region of *SGRAN_1165* |
| **Sgran1165_5** | TATAGGATCCGAGGCATAGGGACCCTCA | PCR of downstream region of *SGRAN_1165* |
| **Sgran1165_6** | TATACCCGGGCCTGCACGGGAATATGGTT | PCR of downstream region of *SGRAN_1165* |
| **KmFw-pk18** | GATTGAACAAGATGGATTGC | PCR to check first integration of pMPO1412 derivative plasmids |
| **KmRev-pk18** | CGTCAAGAAGGCGATAGAAGG | PCR to check first integration of pMPO1412 derivative plasmids |
| **1165 impact fw** | atggcaaagcggcccgg | PCR to clone *SGRAN_1165* gene in the pTYB21 vector |
| **1165 Impact Rv** | ATATTTGGATCCtatgcctcgcgccac | PCR to clone *SGRAN_1165* gene in the pTYB21 vector |
| **3483 Impact Fw 2** | caacagcgccgcctcgg | PCR to clone *SGRAN_3483* gene in the pTYB21 vector |
| **3483 Impact Rv** | ATATTTGGATCCtcagccatggaccgcctg | PCR to clone *SGRAN_3483* gene in the pTYB21 vector |
| **phyR2-FLAG-1** | CATGATTACGAATTCGAGCTCGCTTTGCCATGAATCCGGTTC | C-terminal 3×FLAG tagging of PhyR2 |

| | | |
|---|---|---|
| **phyR2-FLAG-2** | TTAATCATGATCCTTGTAGTCGATGTCATGAT CTTTATAATCACCGTCATGGTCTTTGTAGTC G GCTGCGAGTTGCGGCGTG | C-terminal 3×FLAG tagging of PhyR2 |
| **phyR2-FLAG-3** | ACTACAAGGATCATGATTAATAGGACGGATT GCCATTC | C-terminal 3×FLAG tagging of PhyR2 |
| **phyR2-FLAG-4** | TGCCTGCAGGTCGACTCTAGAGGCTAAGTA ACGACGGAATG | C-terminal 3×FLAG tagging of PhyR2 |

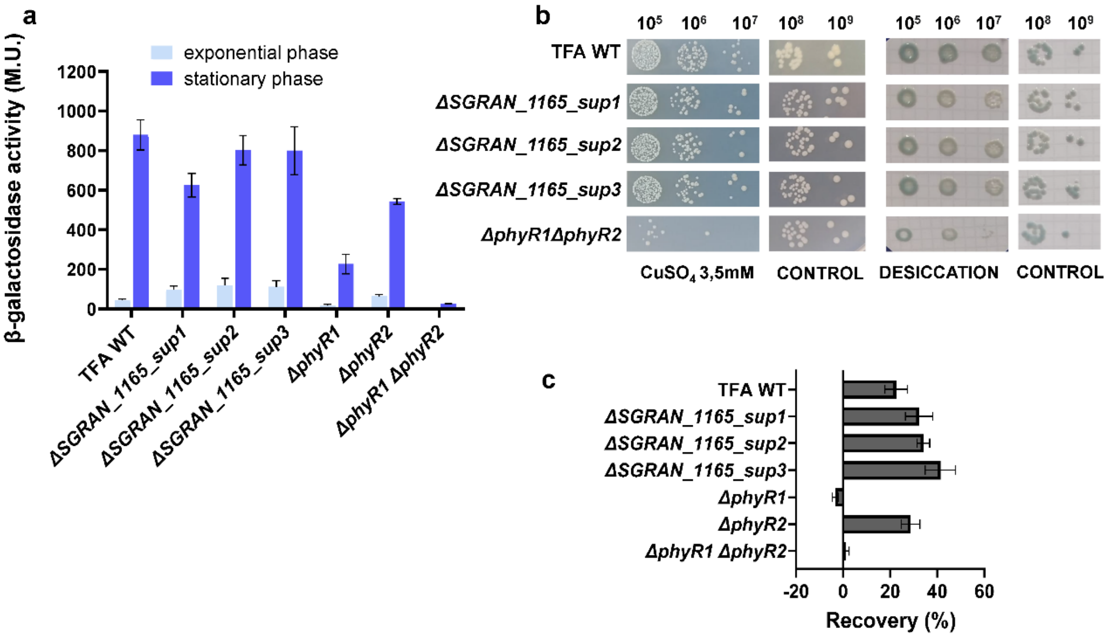


**Figure S1. GSR activity and stress phenotypes of *ΔSGRAN_1165* suppressor mutants.**

GSR-dependent expression and stress phenotypes of three independently isolated *ΔSGRAN_1165 suppressor* mutants, compared with wild-type TFA and the *ΔphyR1ΔphyR2* double mutant.

(a) GSR-dependent expression was assessed during exponential growth (10 h) and stationary phase (48 h) in MM minimal medium with 8 mM β-hydroxybutyrate as the carbon source. Strains carrying a *nepR2::lacZ* translational fusion were used, and GSR-dependent expression was determined by β-galactosidase assays. (b) Heavy-metal stress sensitivity was assessed by spotting serial dilutions onto MML agar supplemented with 3.5 mM $CuSO_4$. Desiccation resistance was assessed by spotting serial dilutions onto filters, air-drying them for 20 h, and subsequently placing the filters onto MML agar supplemented with 0.002% bromophenol blue. Filters transferred directly onto the plates without air-drying were used as controls. (c) Recovery from oxidative shock was assessed by comparing the $OD_{600}$ of cultures exposed to 10 mM $H_2O_2$ with that of untreated controls 3 h after treatment.
